# Cell Cycle Regulation has Shaped Budding Yeast Replication Origin Structure and Function

**DOI:** 10.1101/2024.01.10.575016

**Authors:** Chew Theng Lim, Thomas C.R. Miller, Kang Wei Tan, Saurabh Talele, Anne Early, Philip East, Humberto Sánchez, Nynke H. Dekker, Alessandro Costa, John F.X. Diffley

**Affiliations:** Chromosome Replication Laboratory; Macromolecular Machines Laboratory; Bioinformatics & Biostatistics The Francis Crick Institute, 1 Midland Road, London, NW1 1AT; Department of Bionanoscience, Kavli Institute of Nanoscience, Delft University of Technology, Delft, The Netherlands; DNRF Center for Chromosome Stability, Department of Cellular and Molecular Medicine, University of Copenhagen, Copenhagen, Denmark

## Abstract

Eukaryotic DNA replication initiates from multiple genomic loci known as origins. At budding yeast origins like ARS1, a double hexamer (DH) of the MCM replicative helicase is assembled by Origin Recognition Complex (ORC), Cdc6 and Cdt1 via sequential hexamer loading from two opposed ORC binding sites. Cyclin Dependent Kinase (CDK) inhibits DH assembly, which prevents re-replication by restricting helicase loading to G1 phase. Here we show that an intrinsically disordered region (IDR) in the Orc2 subunit promotes interaction between ORC and the first loaded, closed-ring MCM hexamer (the MO intermediate); CDK phosphorylation of this IDR blocks MO formation and DH assembly. We show that MO functions by stabilising ORC at the lower affinity binding sites required for second hexamer loading. Origins comprising two high affinity ORC sites can assemble DH efficiently without MO by independently loading single hexamers; these origins escape CDK inhibition *in vitro* and *in vivo*. Our work reveals mechanistic plasticity in MCM loading with implications for understanding how CDK regulation has shaped yeast origin evolution and how natural origins might escape cell cycle regulation. We also identify key steps common to loading pathways, with implications for understanding how MCM is loaded in other eukaryotes.

## Introduction

The initiation of eukaryotic DNA replication is separated into two temporally distinct steps that ensures no region of the genome is replicated more than once^1–3^. During G1 phase, a pair of hexameric MCM helicases is loaded at each replication origin around double stranded DNA in the form of a head-to-head MCM DH. During S-phase, each DH is converted into two active CMG (<u>C</u>dc45-<u>M</u>CM-<u>G</u>INS) helicases, thereby initiating bidirectional DNA replication.

Budding yeast DNA replication origins generally contain a high affinity ORC binding site flanked by one or more low affinity sites in the opposite orientation^4–6^. At ARS1 (Fig. 1a), the bipartite high affinity site comprises the A and B1 elements (A/B1) and a lower affinity site corresponds to the B2 element^6–8^. Both the A and B2 elements contain the <u>e</u>xtended <u>A</u>RS <u>c</u>onsensus <u>s</u>equence (EACS): the A element is a closer match to the EACS than B2. DH assembly at ARS1 occurs via sequential loading of two MCM hexamers (outlined in Fig. 1b)^5,9,10^. After loading of the first hexamer (steps i-iii), ORC releases from A/B1. ORC then binds both B2 and the N-terminal domain of the first loaded MCM, generating the <u>M</u>CM-<u>O</u>RC or ‘MO’ complex (iv)^11^. The same ORC that was bound to A/B1 can ‘flip’ and bind B2 (Ref ^12^); alternatively, a second ORC molecule can bind B2 and engage the first recruited MCM (Ref ^5^). In MO, the MCM ring is closed and MCM subunits are primarily in a post-hydrolysis (ADP-bound) state, very similar to the DH^11^. Because this second ORC binding event also induces a DNA bend, the face of ORC that promotes initial MCM recruitment and OCCM formation is available in MO to receive the second MCM-Cdt1 without steric interference (v). MO is thus poised to be a key intermediate in DH assembly (vi); nonetheless, the exact mechanism of MO formation, its function and its importance in DH assembly remain unclear. Moreover, whether all DH assembly proceeds via MO is unknown.

**Figure 1.**
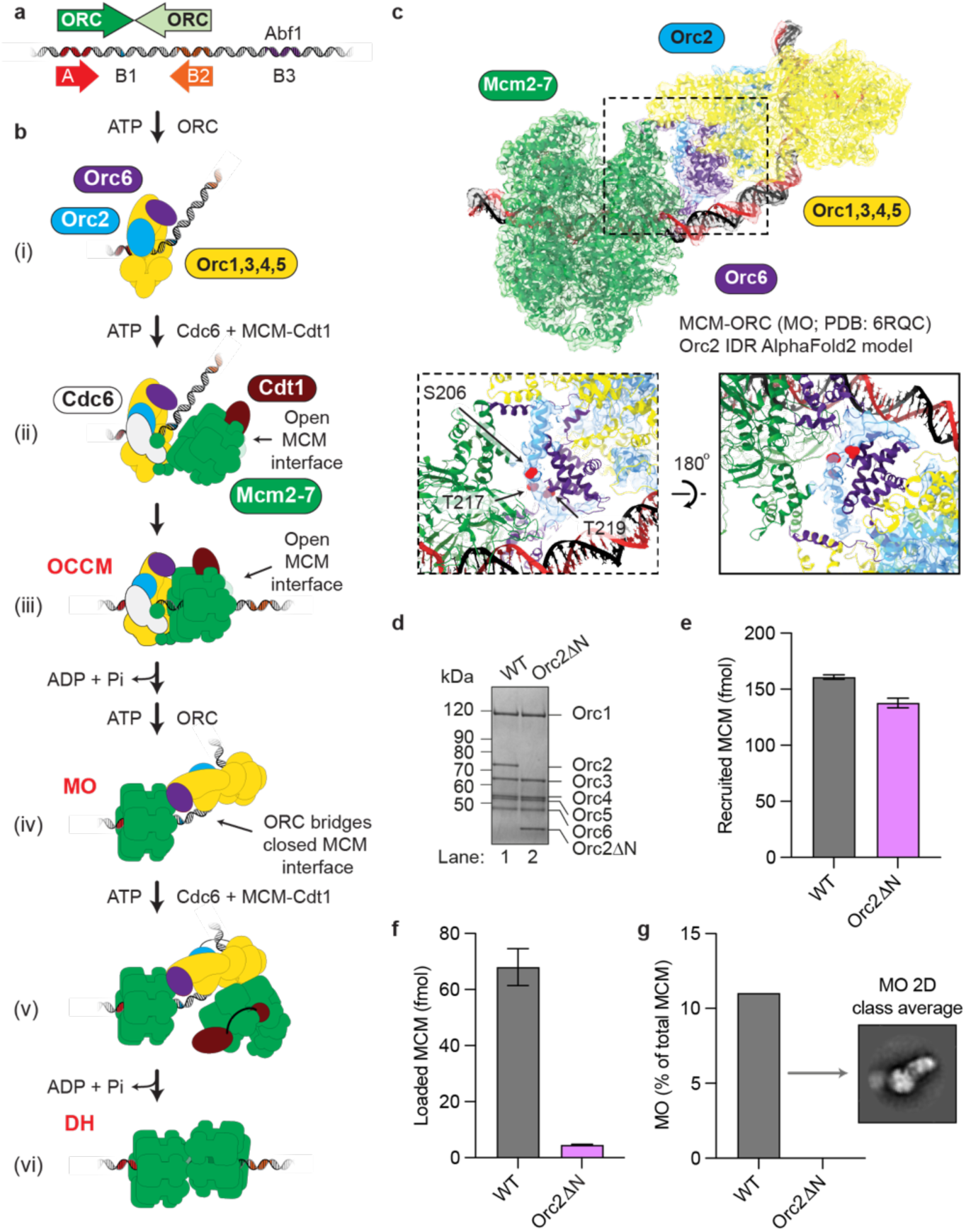
The Orc2 IDR is a key element of the MCM-ORC interface. **a.** Architecture of the canonical ARS1 origin of replication, with high-(dark green) and low-affinity (light green) ORC binding sites. **b.** Schematic of the MCM-DH formation pathway. **c.** Cryo-EM structure of the MCM-ORC intermediate (PDB: 6RQC, EMDB: 498011) including the Orc2 IDR (aas 190-231) modelled by AlphaFold213. The Orc2 IDR, including CDK phosphorylation sites S206, T217 and T219, occupies previously unassigned EM density at the center of the MCM-ORC interface. The Orc2 EM density has been isolated from a density modified (EMready62) map of ORC obtained from the multibody refinement of the MO11. **d.** Orc2ΔN is stably incorporated into the ORC complex. **e.** DNA-pulldown assays show Orc2ΔN recruits luciferase-tagged MCM to ARS1 in ATPψS with comparable efficiency to wild type (WT) ORC. **f.** Orc2ΔN is compromised in MCM loading on ARS1 in ATP. The data in (e) and (f) is plotted as mean ± SD. **g.** MCM loading assays visualized by negative stain EM show that Orc2ΔN fails to form MO on ARS1.

## Results

### MO is crucial for MCM loading at ARS1

We first sought to understand the importance and role of MO in DH assembly by identifying an ORC mutant that cannot form MO. We have previously shown that an N-terminal deletion of the Orc6 subunit of ORC (Orc6Δ119) reduced both MO and DH formation by approximately 50% consistent with the idea that MO is a precursor of DH^11^; however, this was a relatively modest effect and we wanted a mutant with more complete loss of MO formation. Orc2 contains a long (235aa) N-terminal extension that is predicted by AlphaFold^13^ to be intrinsically disordered. The Orc2 IDR has not been resolved in any ORC structure determined experimentally, including MO. However, a region of the Orc2 IDR was tentatively assigned as wrapping around Orc6 in a cryo-EM density map of DNA-bound ORC^14^, albeit at a resolution that prevented assignment of the amino acid sequence and inclusion of this region in the deposited PDB (5ZR1-Ref ^14^). Using AlphaFold2 multimer^15^, we identified a region of the Orc2 IDR docking on the C-terminal domain of Orc6 with high confidence (Fig. 1c;, Extended Data Fig. 1), consistent with the density seen by Li *et al*. Notably, overlaying our AlphaFold prediction with our previously determined structure of the MO intermediate places the region of the Orc2 N-terminal IDR (aas 190-231) into a contiguous stretch of density at the core of the MCM-ORC interface, suggesting that it may play a key role for MO formation (Fig. 1c). Deletion of this IDR from Orc2 did not affect the ability of Orc2 to form a stable complex with the other ORC subunits (Fig. 1d). ORC complexes containing this deletion (Orc2ΔN) recruited MCM normally in ATPγS (Fig. 1e). However, Orc2ΔN loaded MCM at a level only 7% that of wild type ORC (Fig. 1f). To visualise the effect of Orc2 IDR truncation on DH formation directly, we performed MCM loading assays on a 454bp linear DNA template containing ARS1 flanked by nucleosomes and imaged the reactions by negative stain EM. Whilst MO was readily detectable with wild type ORC in ATP, consistent with previous work^11^, MO was not detected in reactions with Orc2ΔN (Fig. 1g). Thus Orc2ΔN can efficiently recruit MCM to DNA but does not form the MO intermediate, and thus cannot form DHs efficiently. These experiments define a role for Orc2 IDR as a key element of the MO intermediate and provide stronger evidence that MO is a critical intermediate in the assembly of the DH at ARS1.

### MO is Required to Stabilise ORC at Weak Binding Sites

The two key ORC binding sites at ARS1 (A/B1 and B2) are so close together that ORC cannot bind both sites simultaneously^6^. Thus, MCM loading must occur in two sequential ORC binding steps. In Fig. 2a we arranged budding yeast origins according to the distance between the best match to the EACS (A domain) and the best predicted secondary ORC site in the opposite orientation (B2-like domain). In roughly 1/3 of yeast origins, the two ORC binding sites are as close or closer together than A/B1 and B2 in ARS1 (Fig. 2a). Using synthetic origins with two high affinity ORC sites (perfect EACS/B1) in which distances are measured from the inside edges of the two B1 elements, 10bp corresponds to the distance between A/B1 and B2 in ARS1, and this construct, like ARS1, can only bind a single ORC molecule (Extended Data Fig. 2a lanes 5 and 6). Moving binding sites just two bp further apart, however, allows two ORC molecules to bind simultaneously (Extended Data Fig. 2a lanes 7 and 8). Thus, roughly 2/3 of yeast origins are predicted to allow simultaneous binding of two ORC molecules (Fig. 2a). In approximately 20% of yeast origins, the two predicted ORC binding sites are far enough apart to fit a single MCM hexamer between them and in 23% they are far enough apart to fit a DH between them. However, in only 15% of origins, the weaker ORC binding site is a better EACS match than ARS1 B2 (Extended Data Fig. 2b). Thus, asymmetry in ORC binding affinities appears to be a general property of yeast origins regardless of whether or not the two sites overlap. The MCM loading mechanism has primarily been studied using ARS1; it remains unclear whether the mechanism of MCM loading is the same at all origins or whether other mechanisms may be important at origins with different ORC binding affinities or wider spacing between ORC binding sites.

**Figure 2.**
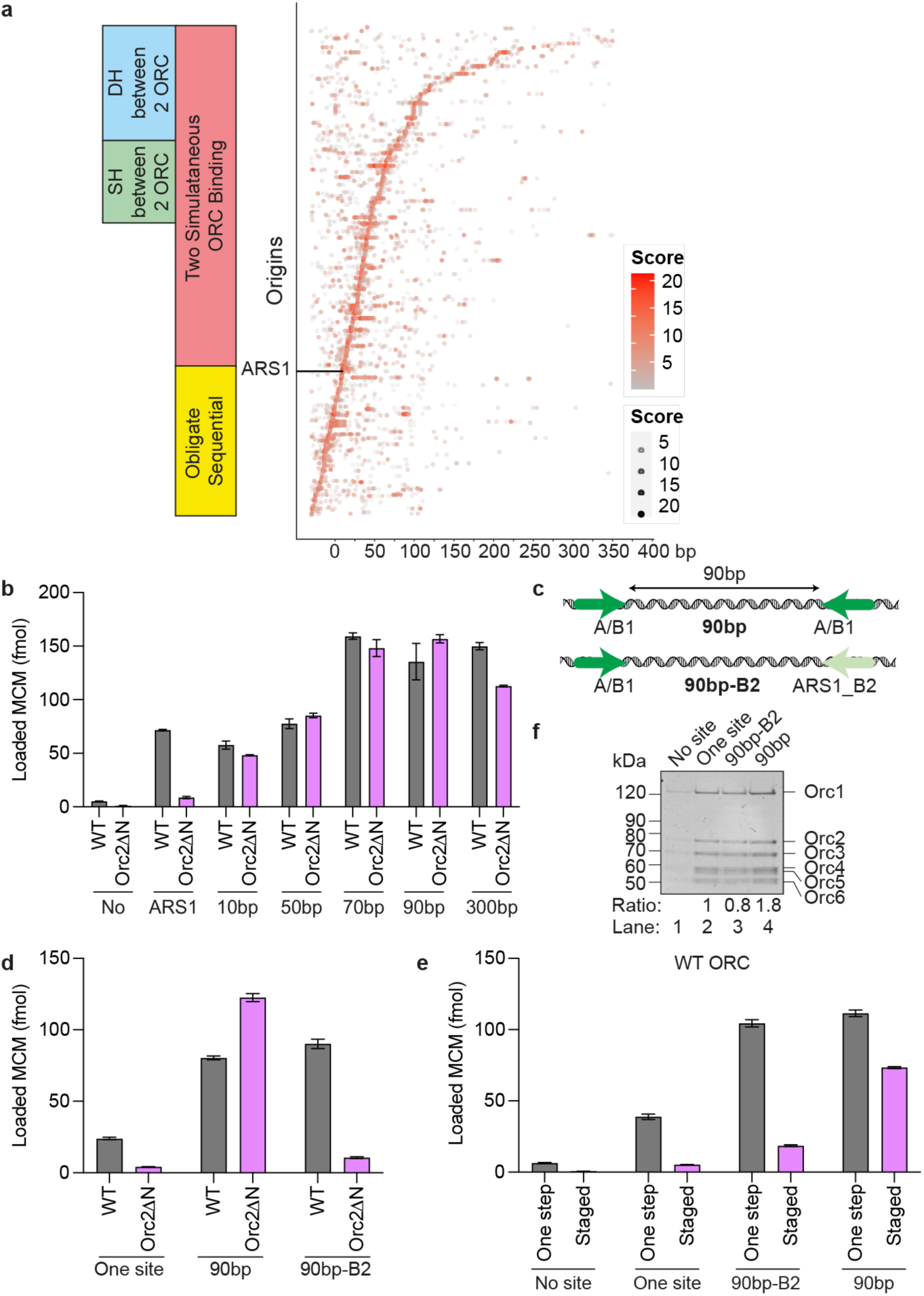
The MO promotes MCM loading by stabilizing ORC at low-affinity binding sites. **a.** Matches to the EACS were identified using a position weight matrix (PWM) as previously described5. Origins are arranged according to the distance between the centre of the best match to the EACS and the centre of the best B2-like EACS by PWM in the opposite orientation. PWM scores indicated by size/colour of dots as indicated in the figure. **b.** Orc2ΔN loads MCMs onto origins containing two high-affinity inverted ORC binding sites spaced ≥10 bp apart with comparable affinity to WT ORC. **c.** Schematic of synthetic origins containing inverted high affinity ORC binding sites spaced 90bp apart (top), or with one high affinity site replaced by the ARS1 B2 element (bottom). **d.** MCM loading on 90bp origins by Orc2ΔN is dependent on two high affinity ORC binding sites. **e.** MCM loading by WT ORC shows two DNA-bound ORCs load MCM more efficiently than one ORC in staged reaction. **f.** ORC is retained on 90bp and 90bp-B2 origins after washing. The binding ratio is normalised to the one site template, showing 90bp symmetrical origins retain approximately twice as much ORC as 90bp-B2 origins.

To begin to address this, we tested the ability of Orc2ΔN, which cannot form MO, to load MCM on a series of synthetic origins with two high affinity ORC binding sites in the opposite orientation and placed at distances from 10 to 300 base pairs apart. As shown previously^5^, wild type ORC can load MCM efficiently on this series of origins, with a peak of MCM loading at 70-90bp but with efficient loading even at 300bp. Orc2ΔN cannot load MCM efficiently at ARS1, but surprisingly, loads MCM at all distances from 10 to 300bp as efficiently as wild type ORC (Fig. 2b). This suggests that MO is not required when MCM is loaded from two high affinity sites, regardless of whether those sites effectively overlap (10bp) or are separated by a relatively large distance (300bp). To test this directly, we changed one high affinity EACS/B1 site in the most efficient synthetic template (90bp) to the B2 element from ARS1, a lower affinity ORC site (Fig. 2c). As shown in Fig. 2d, wild type ORC loads MCM as efficiently on this template as it does on the parent template with two high affinity sites, demonstrating that weak ORC sites can be effectively used even when the two ORC sites do not overlap. MCM loading with Orc2ΔN on the 90bp template was at least as efficient as wild type, but MCM loading on the 90bp-B2 construct was greatly reduced.

From these experiments we conclude that the MO intermediate is critical for MCM loading when one of the ORC binding sites is lower affinity, and, therefore, that the essential role of MO in DH assembly is to stabilise ORC binding at weak sites after first hexamer loading. Furthermore, MO is still essential when the two ORC sites are 90bp apart indicating that MO is not just required when ORC sites overlap, as in ARS1. However, MO is not required when MCM is loaded from two high affinity ORC binding sites.

Because the EACS/B1 sequences in both 90bp and 90bp-B2 are high affinity ORC binding sites, ORC binding to these sequences is stable to stringent (80 mM NaCl) washing whilst binding to B2 is not^5^, which gives us an opportunity to examine the efficiency of the ‘one ORC’ loading via ORC flipping vs the ‘two ORC’ mechanisms where a second ORC molecule loads the second hexamer. To test this, reactions were ‘staged’: near-saturating amount of wild type ORC was pre-bound to either the 90bp or the 90bp-B2 origin (Extended Data Fig. 2c), then free ORC along with B2-bound ORC was washed off (Extended Data Fig. 2d). We then added Cdc6 and MCM-Cdt1 and examined MCM loading. As shown in Fig. 2e, in staged reactions the 90bp template loaded approximately 4 times more MCM than the 90bp-B2 template (pink bars) despite the fact that both templates work equally well in a one-step reaction (grey bars) and ORC was retained on both templates after washing (Fig. 2f). This result is consistent with the idea that a single ORC can load a double hexamer from a single high affinity site via ORC flipping and MO formation but also shows that this pathway is less efficient than the pathway in which each MCM single hexamer is loaded by separate ORC molecules at two high affinity sites. Moreover, the difference between the one-step and staged reaction with 90bp-B2 also argues that the single ORC pathway is less efficient than the two ORC pathway even when loading works through MO. Therefore, we can distinguish three separate ways MCM can be loaded: one ORC via MO, two ORC via MO and two ORC MO-independent.

### CDK phosphorylation of Orc2 blocks MO formation

CDK promotes Cdc6 proteolysis^16–19^ and MCM-Cdt1 nuclear export^20–24^ and also blocks ORC’s ability to assemble DHs *in vitro*^25–27^;. The region in the Orc2 IDR predicted to form part of the MCM-ORC interface in MO (Fig. 1c and yellow box in Extended Data Fig. 3a) is moderately conserved amongst closely related budding yeasts, which all contain multiple potential CDK phosphorylation sites near the predicted interface with MCM in MO (Fig. 1c red dots). We therefore hypothesised that CDK phosphorylation of Orc2 might inhibit MCM loading by interfering with MO formation. Phosphorylation of wild type ORC with S phase CDK (Clb5-Cdc28-Cks1) caused a reduction in mobility of both the Orc2 and Orc6 subunits in SDS PAGE (Fig. 3a); this CDK phosphorylated ORC was very inefficient in DH formation at ARS1 (Fig.3b), consistent with previous work^25–27^. Similar to Orc2ΔN, however, phosphorylated ORC loaded MCM almost as well as unphosphorylated ORC on the 90bp synthetic origin (Fig. 3b). To ensure this loaded MCM was functional, we tested its ability to support DNA replication *in vitro* with purified proteins. Whilst pre-phosphorylated ORC did not support replication from an ARS1-containing template (compare lanes 5 and 6, Fig.3c), it supported replication from a template containing the 90bp origin almost as well as unphosphorylated ORC (lanes 3 and 4, Fig. 3c and Extended Data Fig. 4a) indicating that the DH formed is functional. Consistent with this, a 2.8 Å resolution cryo-EM structure of DH assembled with phosphorylated ORC is virtually identical to DH assembled with unphosphorylated ORC (Extended Data Fig. 5a). To test whether this 90bp origin could also bypass CDK regulation of ORC *in vivo*, we used a yeast strain in which CDK regulation of MCM was constitutively eliminated by fusing an unregulated nuclear localisation sequence (SV40-TAg NLS) to Mcm7, and CDK regulation of Cdc6 was conditionally deregulated with a copy of a stable version of Cdc6 (Cdc6ΔNT) under a galactose-inducible promoter. When both Cdc6 and MCM are deregulated, cells rely entirely on wild type ORC phosphorylation to prevent re-replication^28^. We integrated a series of origins in place of ARS419, then arrested cells in G2/M, expressed stable Cdc6 and examined copy number after 3 hrs. As shown in Fig. 3d, deregulation of Cdc6 and MCM did not lead to any increase in DNA copy number of ARS1; ARS317, which had been previously shown to re-replicate under similar conditions^29^, showed elevated copy number, but the 90bp origin showed the largest increase in copy number after Cdc6 deregulation consistent with the idea that this synthetic origin bypasses CDK regulation of ORC both *in vitro* and *in vivo*.

**Figure 3.**
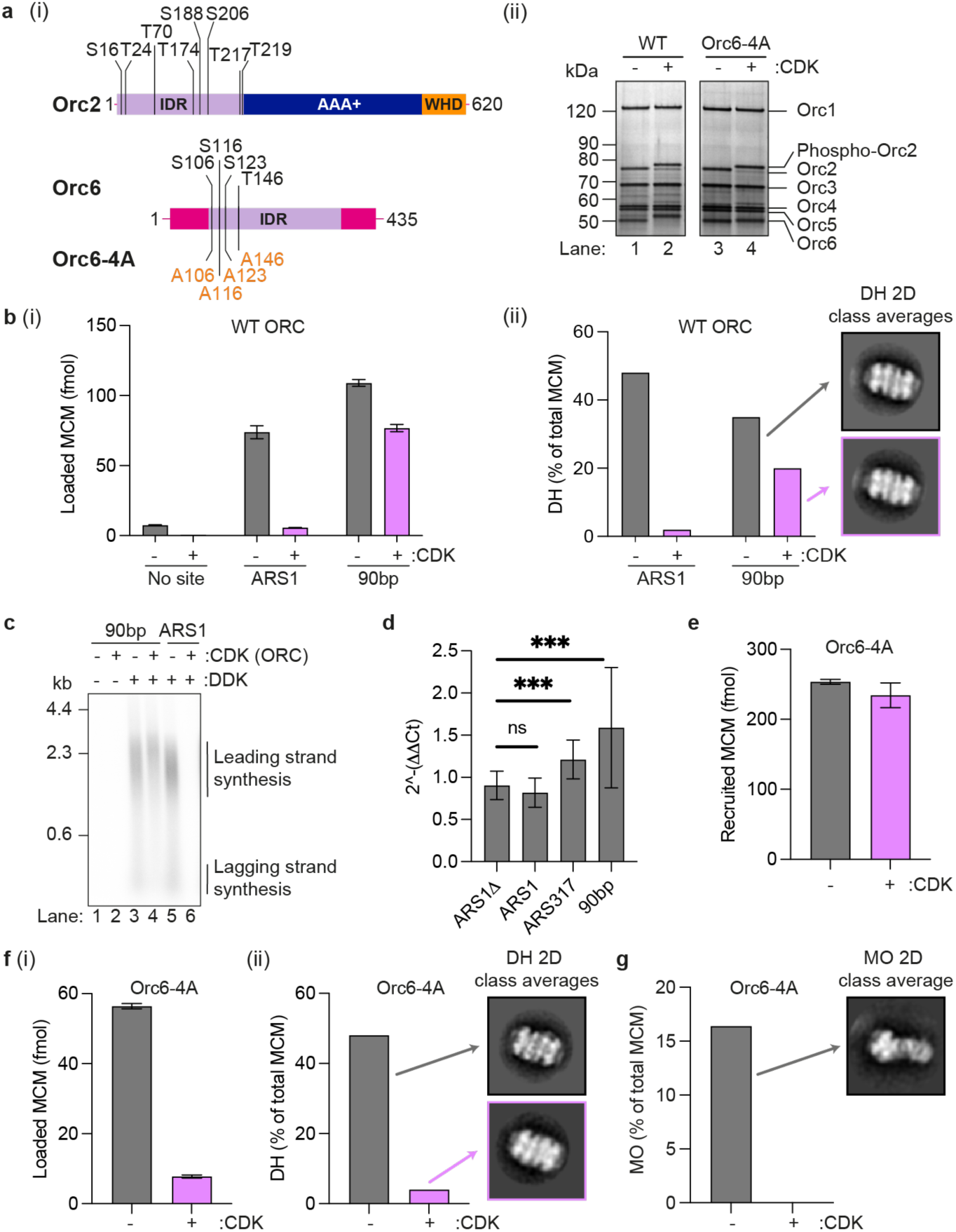
CDK phosphorylation of Orc2 inhibits the formation of MO. **a.** (i) Potential CDK phosphorylation sites (consensus sequences (^S^/TPX^K^/R)) in Orc2 and Orc6. (ii) An Orc6 mutant with four CDK target sites mutated to alanine (Orc6-4A) can be stably incorporated into the ORC complex but not phosphorylated. CDK phosphorylation of Orc2 is comparable to WT. **b.** MCM loading on ARS1, but not 90bp origins, is highly sensitive to CDK phosphorylation ORC. **c.** MCMs loaded onto 90bp origins by phosphorylated ORC are replication competent. **d.** In yeast cells dependent on ORC phosphorylation to prevent re-replication, 90bp spaced origins inserted into the yeast genome re-replicate causing an increase in copy number, as detected by qPCR. ORC phosphorylated on Orc2 by CDK (Orc6-4A mutant) efficiently recruits MCMs to ARS1 origins **(e)** but is inhibited in MCM loading **(f)**. **g.** ORC phosphorylated on Orc2 by CDK (Orc6-4A mutant) fails to form MO.

We next sought to determine which step in DH assembly was blocked by Orc2 phosphorylation. To examine the effects of Orc2 phosphorylation alone, we mutated four Ser/Thr residues in the CDK consensus sequences (^S^/_T_PX^K^/_R_) in Orc6 to alanine to generate Orc6-4A. As shown in Fig. 3a, the phosphorylation-dependent shift of Orc6 was greatly reduced in Orc6-4A, but Orc2 was still shifted by CDK phosphorylation. On ARS1, CDK phosphorylation of Orc2 did not affect MCM recruitment in ATPγS (Fig. 3e). However, Orc2 phosphorylation led to a substantial reduction in DH formation in both DNA-pulldown (Fig. 3fi) and EM-based (Fig. 3fii) assays and a complete inhibition of MO formation (Fig. 3g), indicating that phosphorylation of Orc2 alone is sufficient to inhibit DH formation. Thus, CDK phosphorylation of Orc2, like deletion of the Orc2 IDR (Fig. 1), blocks DH assembly before MO formation. Whilst CDK phosphorylation of Orc2 blocks MO formation and DH formation on the ARS1 origin, it does not block DH formation on the 90bp origin (Fig 3b), consistent with the results in the previous section showing that MO is not required when origins contain two high affinity binding sites (Fig. 2b and d).

**Figure 4.**
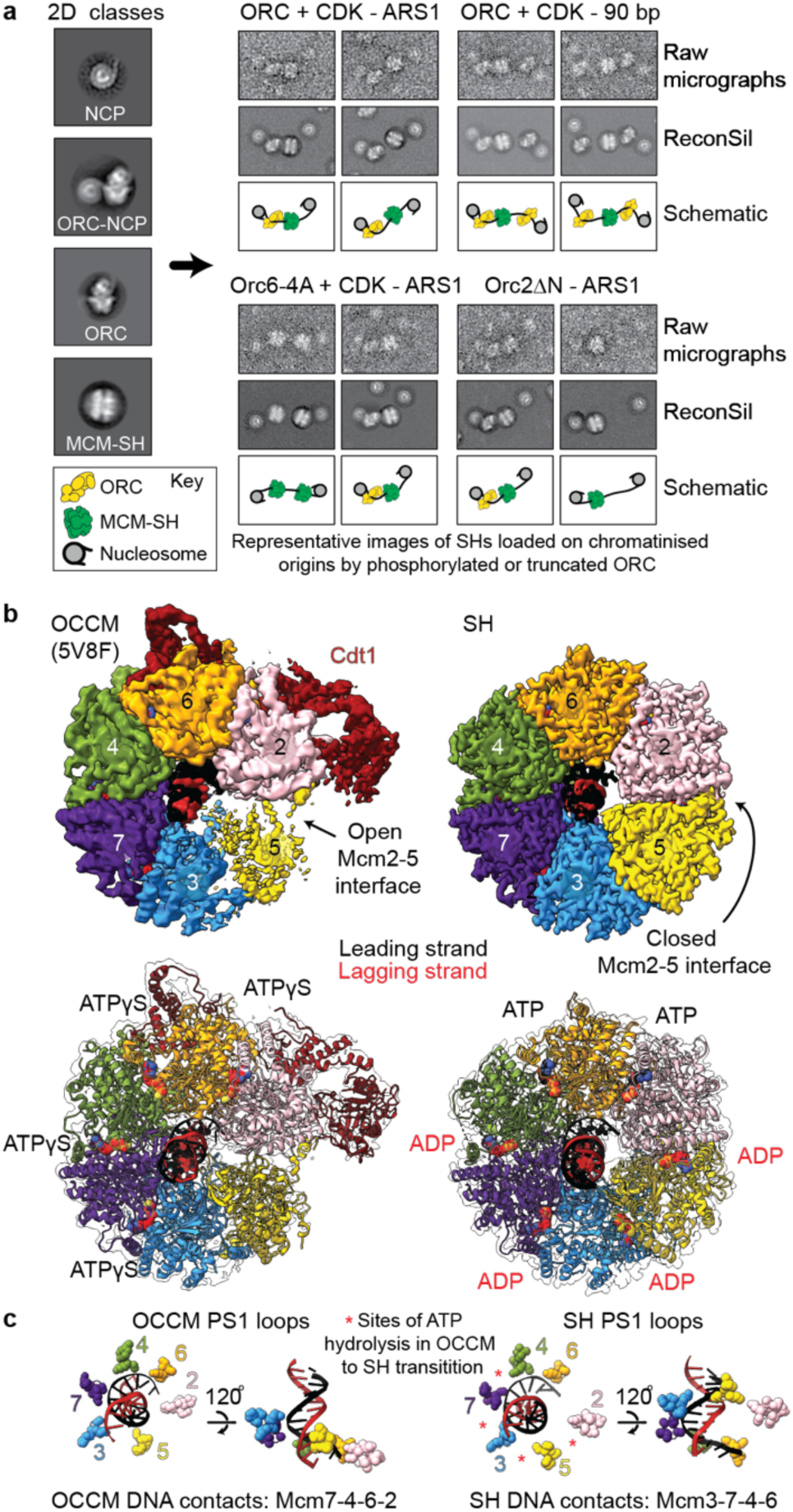
Structure of the MCM SH a. Negative stain EM in silico reconstitution (ReconSil) of MCM loading reactions performed with phosphorylated or truncated Orc2. Left, conventional EM 2D class averages. Right, representative images of chromatinised origins bound by single MCM helicases and/or DNA-bound ORCs flanked by nucleosomes, shown as raw micrographs (top rows), after ReconSil (middle rows) or as schematics (bottom rows). **b.** Cryo-EM structures and atomic models for OCCM (left; 5V8F [Ref 65]) and a SH (right; this study). The EM density map displayed for the SH has been density modified using EMReady62. **c.** ATP hydrolysis in the Mcm4-7, 7-3, 3-5 and 5-2 ATPase sites during SH loading is associated with a reconfiguration of the MCM PS1 loops into a staircase configuration that engages the leading strand template DNA backbone.

### Structure of the MCM single hexamer

Formation of MO requires Orc2 and Orc6 to bridge the N-terminal interface between the Mcm2 and Mcm5 subunits that form MCM’s DNA entry ‘gate’. However, it is currently unknown whether closure of the Mcm2-5 interface precedes ORC binding in the MO or whether ORC binding closes the Mcm2-5 gate to load the first MCM hexamer. Phosphorylated Orc2 (Orc6-4A) recruits MCM normally but does not form MO suggesting that Orc2 phosphorylation either blocks a step in the loading of the SH, for example OCCM disassembly, and/or directly prevents ORC binding to a loaded SH during the formation of MO. To investigate this, we performed single particle reconstitution in silico (RECONSIL)^11^ experiments to visualise entire chromatinised origins of replication during MCM loading reactions containing phosphorylated or truncated Orc2. RECONSIL experiments enabled us to identify SHs apparently trapped on DNA between nucleosomes (Fig. 4a). Thus, the Orc2 IDR seems not to be required for OCCM assembly or disassembly to release SH onto DNA but is required for MO-formation, which can be inhibited by CDK phosphorylation of the Orc2 IDR.

To confirm this, we used cryo-EM to visualise the helicase loading reaction with CDK-phosphorylated Orc2. The 3.4 Å resolution structure (3.1 Å after density modification) shows a single MCM ring encircling duplex DNA (Fig. 4b, Extended Data Fig. 5). Unlike the MCM in the OCCM structure, the SH contains a completely closed Mcm2-5 interface identical to that seen in the DH (Fig. 4b, Extended Data Fig. 5b). Thus, phosphorylation of ORC2 blocks DH formation after a SH has been fully closed around DNA, implying that ring closure precedes and is not a consequence of N-terminal ORC binding that results in MO formation. Comparison with MCM in the OCCM complex reveals that the DNA grip in the helicase ring changes after closure of the Mcm2-5 gate (from Mcm7-4-6-2 to Mcm3-7-4-6 duplex engagement), likely as a consequence of ATP hydrolysis (Fig. 4b). Nucleotide occupancy changes from ATPγS in the 3-7, 7-4, 4-6 and 6-2 interfaces in OCCM, to ATP in the 4-6 and 6-2 interfaces and ADP in the 2-5, 5-3, 3-7 and 7-4 interfaces in SH. This change in nucleotide occupancy indicates a minimum of 4 ATP hydrolysis events that may drive the re-orientation of the ATPase pore loops, with pre-sensor 1 (PS1)rranged in a staircase configuration that follows the helical rise of the leading-strand template (Fig. 4c), as observed in the active CMG helicase (Extended Data Fig. 5c).

### Mechanism of MO-independent DH assembly

To examine the properties of the loaded SH, we assembled MCM with Orc2ΔN on either a one site template, where only SH assembles, or a template containing two ORC binding sites separated by 90bp where DH efficiently forms; we then challenged the loaded MCM products with buffers containing different salt concentrations. As shown in Fig. 5a, MCM loaded on both templates was resistant to salt concentrations up to 250 mM NaCl; virtually all of the MCM was removed from the one site templates at 500 and 1000 mM NaCl whilst about 40% of the MCM remained on the 90bp template after 500 and 1000 mM NaCl washes. These experiments indicate that SH is stable up to ∼ 250 mM NaCl, but is efficiently removed at salt concentrations of 500 mM NaCl and above. The DH is stable up to 2 M NaCl^30^, so the fraction of MCM removed by 500 mM NaCl from the 90bp origin likely represents SH, with the MCM remaining at 500 and 1000 mM NaCl being DH. This suggests that a substantial fraction of the MCM loaded even on this very efficient origin are SHs that have not matured into DHs. This is consistent with single molecule experiments showing that SHs are readily detected during normal MCM loading reactions^31^.

**Figure 5.**
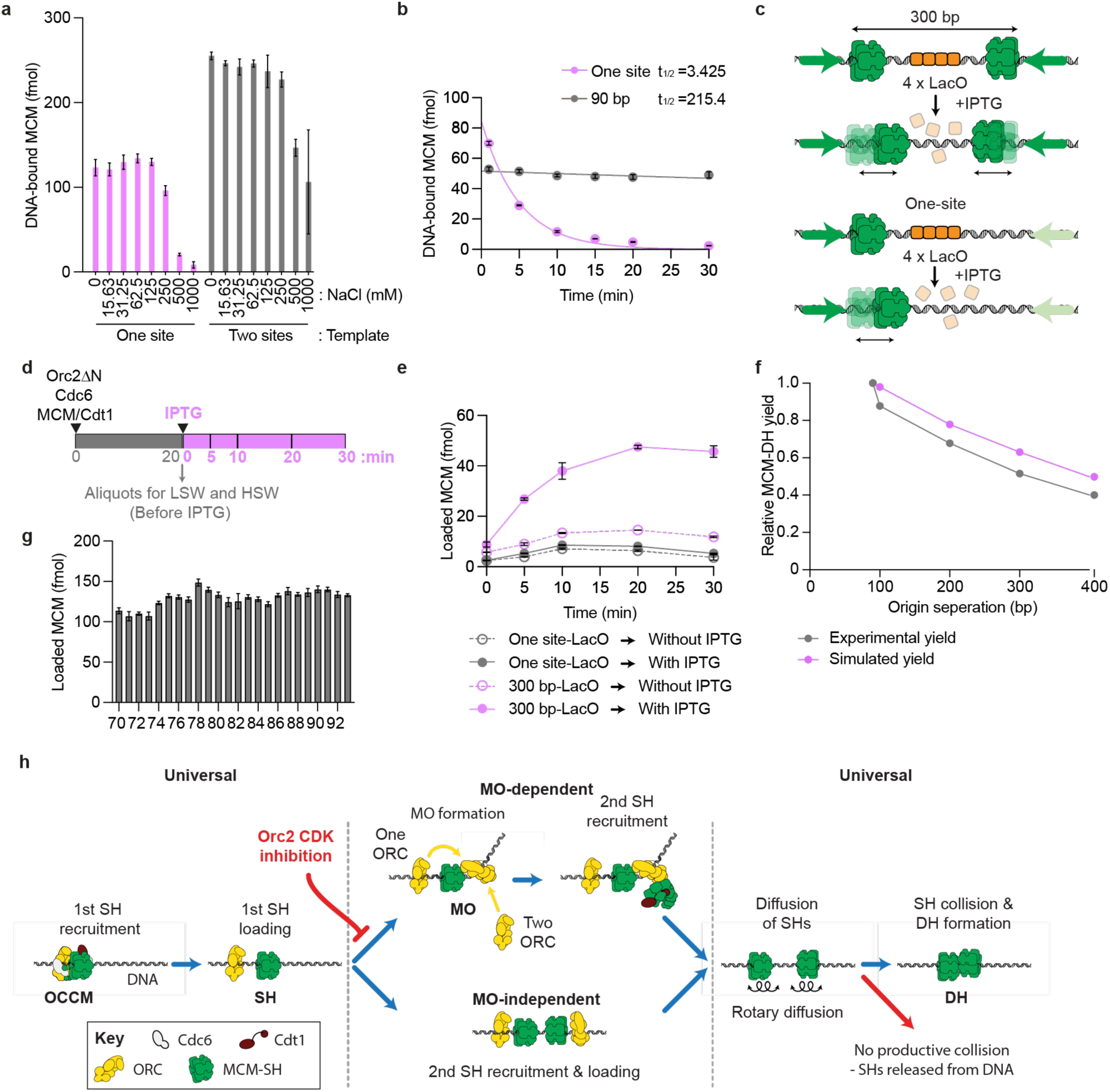
Mechanism of MO-independent DH formation. SHs have a lower salt stability **(a)** and shorter half-life **(b)** on DNA than DHs. Schematics illustrating origins **(c)** and assay **(d)** used for characterising MO-independent DH formation via collision of two independently loaded SHs **(e)**. **f.** Efficiency of DH formation decreases with increase in separation between ORC binding sites and can be modelled as the meeting probability of two diffusing SHs loaded at ORC binding sites at a given separation in the experimental time (20 minutes). **g.** MCM loading is insensitive to the separation distance between two ORC binding sites (between 70-93 bp) and relative orientation of recruited ORCs with respect to each other along the DNA helix. **h.** Proposed model for MCM loading via MO-dependent and independent pathways. Both pathways converge on a ‘universal’ mechanism that requires loading of two SHs that must collide in a head-to-head orientation to form a DH.

To determine the off rate of SH under MCM loading conditions (low salt), MCM was loaded as above using Orc2ΔN on a single site sequence, excess protein was removed by low salt wash, and the amount of SH remaining on DNA was then assessed after incubation for the indicated times followed by low salt wash. Fig. 5b shows that SHs were lost from DNA with a half-life of approximately 3.5min. To compare this to the DH, we loaded MCM onto the 90bp template then washed free proteins and SHs off with high salt (1 M NaCl) before taking time points. As shown in Fig. 5b, DHs are stable for the entire duration of the experiment. Thus, the SH is much less stable than the DH on DNA.

To show more directly that two independently loaded SHs can assemble into a DH, we generated a template with two high affinity ORC sites separated by 300bp and 4 lac operators between the sites (Fig. 5c). As a control we also generated a template with a single ORC site and 4 lac operators. We pre-assembled SHs on these templates with Orc2ΔN for 20 minutes (Fig. 5d). As shown in Extended Data Fig. 6a, there is considerable MCM loaded on both templates at this point; however, this MCM is all removed by high salt wash indicating that there are no DHs. DH formation (high salt resistant MCM) was detected at significant levels only on the two ORC binding site template and only after IPTG addition (Fig. 5e) in a time-dependent manner. This strongly supports the idea that two SHs can generate a DH after interaction.

The efficiency of DH assembly decreases as two ORC sites are moved further apart. We reasoned that the further apart two SHs are loaded, the higher the probability that one or the other will fall off before meeting. To model this, we used the observed one dimensional diffusion constant for the DH^32^, together with the off rate determined in Fig. 5b (Extended Data Fig. 9 and 10), and as shown in Fig. 5f, the predicted loading efficiencies are remarkably close to those seen in the experimental data. Diffusion could occur with rotation following the path of the DNA helix, or by diffusion without rotation. For several reasons we favour the idea that movement occurs coupled with rotation tracking the DNA helix. First, the structure of the SH shows a single mode of DNA engagement within the MCM central channel, with individual nucleotides clearly resolved (Extended Data Fig. 5d). Despite the large number of particles (∼400k), further 3D classification failed to detect SHs with an alternate DNA engagement, suggesting that the DNA register remains the same in all imaged SHs, regardless of their relative position on the origin DNA. If SHs diffused without rotation, they would be unlikely to have a highly preferred mode of DNA engagement. Second, without rotation, two SHs would meet in different registers depending on how far apart the loading sites were. One might expect to see evidence of this in loading efficiency; however Fig. 5g shows that loading efficiency is very similar at every distance from 70 to 93bp. Rotation along the DNA helix ensures that the two SHs always meet with the same register; using the first steric clash between hexamers as the end point of rotational diffusion along B-form DNA, this register is very close to the register seen in the DH (Extended Data Movie 1). Taken together, we propose DH assembly occurs by random rotational diffusion and meeting of two SH along the DNA followed by a presently uncharacterised ‘locking’ of the two into a stable DH.

## Discussion

Our results, summarised in Fig. 5h, indicate that MCM can be loaded by at least three distinct pathways in budding yeast. In each pathway, the unifying mechanism is the loading of two SHs with inverted orientations. DH assembly can be coordinated by the MO, which facilitates ORC recruitment to a weak DNA site. Alternatively, DH assembly can be done in the absence of the MO, in which case two independently loaded SHs diffuse and rotate until meeting to form a stable DH. The loading of the SH defines a precise register of DNA engagement that ensures two loaded SHs with inverted orientations will always meet in the same subunit register, regardless of the distance between their loading sites. Our data (Fig. 2b) shows that DH formation is at least as efficient without MO, as with MO, indicating that the SH to DH ‘locking’ mechanism is intrinsic to two correctly oriented and DNA engaged SHs and is independent of the MO.

SHs are considerably less stable on DNA than DHs. We suggest that this is because the Mcm2-5 interaction is intrinsically weak, providing a gate for DNA to exit the central channel, consistent with recent FRET experiments^33^. The Mcm2-5 gates are not aligned in the DH but are located approximately 180° apart, so, even if both gates open at the same time, the DH would remain topologically bound around DNA^34^ which may prevent dissociation of the DH. Alternatively, contacts between the hexamers may stabilise the Mcm2-5 gate in the DH, preventing opening^33^.

Previous work in yeast has shown that a substantial fraction of MCM is released from G1 chromatin with a moderate salt concentration (350 mM NaCl)^35^ suggesting the presence of SH on chromatin. From yeast to humans, the total amount of MCM bound to chromatin far exceeds the number of active replication origins. This ‘MCM paradox’ has been explained at least in part by the existence of dormant origins which are activated under stress conditions^36–39^; our results suggest that, in addition, some of the excess MCM may be in the form of SH, which cannot support replication (e.g. Fig. 3c). Though SHs are relatively short lived, there may be active mechanisms to remove them from chromatin. Further work is required to assess what fraction of chromatin-bound MCM is SH.

Synthetic origins comprising two high affinity ORC binding sites load MCM even more efficiently than ARS1 (e.g. Fig. 2b) and function as autonomously replicating sequences *in vivo*^5^. However, yeast origins generally have a single high affinity site flanked by low affinity sites, indicating that they likely assemble DH primarily via MO. Our work suggests an explanation for this: origins with two high affinity sites cannot be inhibited by CDK phosphorylation of Orc2 (Fig.3 c,d). We propose this arrangement has been selected against during evolution: co-evolution of sequence-specific DNA binding by ORC^40,41^ and CDK regulation of ORC function has resulted in the asymmetry in modern yeast replication origins.

Human Orc2 also contains a long N-terminal IDR^42^ but has no detectable sequence similarity to the yeast IDR. Whether human ORC loads MCM via an MO mechanism remains to be seen. Orc2 from multicellular plants entirely lacks an N-terminal IDR and therefore plants are unlikely to use the same MO-dependent mechanism. Whether plants and metazoans use a mechanism related to mechanism 3 or some other, novel mechanism requires further work. Finally, the recent sequencing of several metamonad genomes has shown that, while all six MCM genes are present, organisms in the genus *Carpediemonas* lack all ORC subunits, Cdc6 and Cdt1 (ref ^43^) indicating additional MCM loading mechanisms remain to be discovered.

## Materials and Methods

### Strains

A full list of primers, plasmids and strains used for strain construction are listed in Supplementary Materials. The YJL3239 strain was a gift from Joachim Li^44^. The ARS317, ARS1, ARS1Δ and synthetic 90bp origins were inserted in ARS419 using the method described in Richardson and Li, 2014^45^ with some modifications. The endogenous ARS317 was deleted and replaced with kanMX6 from pFA6a-3HA-kanMX6 with amplification using CTL61 and CTL62 (yCTL5). URA3 was replaced with hphNT1 from pFA6a-hphNT1 with amplification using CTL98 and CTL99 (yCTL18). URA3 was amplified using pRS306 with primers CTL86 and CTL87 and additionally inserted in ARS419 at position 567 kb in Chromosome IV (yCTL19). The ARS317, ARS1, ARS1Δ and synthetic 90bp origins were amplified from plasmid ARS317-Abf1(pCTL69), ARS1-Abf1 (pCTL68), ARS1Δ (pGC404 in Coster and Diffley, 2017^5^) and 90bp-Abf1 (pCTL67) respectively using CTL160 and CTL161 and counter-selected with 5-FOA, resulting in strains yCTL30, yCTL29, yCTL28 and yCTL33. 1 mg/ml of 5-FOA was added to minimal media containing 1X YNB, 2% glucose, 1X CSM (Complete Supplement Mixture, Formedium, UK) and 80 μg/ml uracil. The ARS317-RIP sequence was identified according to the L4-L12 region in Richardson and Li, 2014^45^.

The sequences for Orc6 phosphorylation mutant Orc6-4A was obtained from Invitrogen GeneArt Synthesis (Thermo Fisher Scientific). This sequence was then used to replace the wild type Orc6 in the plasmid pJF18 as in Frigola *et al.*, 2013^46^ to create strain yCTL6 (Orc6-4A). Endogenous Orc6 was Flag-tagged to allow removal of the endogenous wild-type protein.

Amino acids 2-235 in Orc2 were deleted in the Orc2-N-terminal truncation mutant (Orc2ΔN). This mutant was introduced into pJF19 as in Frigola *et al.* (2013)^46^ to replace the wild type Orc2. The wild-type Orc2 was Flag-tagged so that the endogenous protein could be removed. The resulting strain was named yCTL17.

### Protein expression and purification

In this study, ORC, Cdc6, Luc-MCM/Cdt1, CDK and Sic1 were expressed and purified according to previously described methods in the references Frigola *et al.*, 2013^46^, Coster and Diffley, 2017^5^, Yeeles *et al.*, 2015^47^. The Orc2ΔN mutant was purified identical to wild-type ORC, but with the additional step of using Flag beads after CBP pull-down to remove endogenous wild-type proteins. The same was done for the Orc6 phosphorylation mutants.

### ORC binding assay

ORC binding assays were performed as described in Coster and Diffley, 2017^5^ with some modifications. Briefly, 0.15 pmol DNA of 3 kb PCR substrates with a single biotin (GC227 and GC228) were coupled with 5 μl of Dynabeads M-280 streptavidin (Invitrogen 11205D) in 10 μl of binding buffer (5 mM Tris-HCl, pH 8.0, 0.5 mM EDTA, 1 M NaCl), for 30 mins at 30°C with mixing at 1250 rpm in a 1.5 ml microcentrifuge tube. Beads were washed twice and resuspended in buffer containing 10 mM HEPES-KOH pH7.6 and 1 mM EDTA. The bead-bound DNA substrate was then incubated with 12 nM ORC in 40 μl loading buffer (25 mM HEPES-KOH pH7.6, 100 mM NaOAc, 10 mM MgOAc, 0.02% NP-40, 5% glycerol, 1 mM DTT, 5 mM ATP and 80 mM KCl). The reaction was incubated for 15 mins at 30°C with mixing at 1600 rpm on 96-well plate. Reactions were then washed with low salt wash (LSW) additionally containing 80 mM NaCl twice (45 mM HEPES-KOH pH7.6, 5 mM MgOAc, 300 mM NaOAc, 0.02% NP-40, 10% glycerol, 2 mM CaCl_2,_80 mM NaCl) for 2 mins, 30 °C at 1600 rpm. The washed beads were then washed once with LSW once before resuspending in 12 μl LSW with 600 units of micrococcal nuclease (MNase, NEB M0247S) and incubated for 2 min, 30°C at 1600 rpm. The supernatant from two reactions were collected and separated by Criterion XT Tris Acetate 3-8% (Bio-rad) SDS-PAGE and visualized by silver staining.

### MCM loading

The MCM loading assay was performed as described in Coster and Diffley, 2017^5^. Biotinylated DNA substrate was attached to magnetic beads and then incubated with 12 nM ORC, 20 nM Cdc6, 80 nM Luc-MCM/Cdt1 in 40 μl loading buffer for 20 mins at 30°C with mixing at 1600 rpm. Beads were washed twice with high salt wash (HSW, 45 mM HEPES-KOH pH7.6, 5 mM MgOAc, 0.02% NP-40, 10% glycerol, and 1 M NaCl) for 2 mins and twice with LSW (45 mM HEPES-KOH pH7.6, 5 mM MgOAc, 300 mM NaOAc, 0.02% NP-40, 10% glycerol and 2 mM CaCl_2_) 20 secs. Proteins were released by 30 μl of low salt wash with MNase treatment for 2 mins and transferred to white flat bottom, half area 96-well plates (Corning, 3642). Nano-Glo Luciferase assay substrate (Promega N1120) were diluted 1:50 and 30 μl of diluted substrate was added to the sample. A standard curve was also prepared with 2.5-fold dilution of Luc-MCM (ranging from 0.027 to 256.33 fmol). The luminescence was measured on a PHERAstar plate reader using “LUM Plus” optic module (BMG Labtech) with the read time of one second per well at 12.1 mm focal height. The background (low salt wash only) was subtracted and the amount of Luc-MCM was calculated using a linear regression of standard curve of Luc-MCM. All experiments were done in triplicates.

In the one step reaction in Fig. 2e, 40 nM ORC, 40 nM Cdc6 and 80 nM MCM/Cdt1 were used to saturate the MCM loading activity.

In the staged reaction in Fig. 2e, 40 nM ORC was preincubated with DNA beads for 15 min at 30°C with mixing at 1600 rpm. The reaction was washed twice with LSW+80 mM NaCl buffer described above. Then, 40 nM Cdc6 and 80 nM MCM/Cdt1 were added to the reaction and incubate for another 10 min.

In the inducible roadblock assay, beads were resuspended in 20 μl binding buffer (25 mM HEPES-KOH pH7.6, 100 mM NaOAc, 10 mM MgOAc, 0.02% NP-40, 5% glycerol, 5 mM DTT and 80 mM KCl). Then, 2.4 pmol of LacI (a gift from George Cameron and Hasan Yardimci) was added and incubated for 30 mins at 30°C with mixing at 1250 rpm. The beads were washed twice and resuspended in 5 μl of binding buffer. This mixture was subsequently added to the MCM loading reaction and incubated for 20 min. Following this, 10 mM of IPTG and a 50X molar excess of competitor DNA containing one ORC binding site (EACS.B1 F and EACS.B1 R) were added. The reaction was aliquoted at indicated time points for HSW, followed by LSW, as described above.

### MCM recruitment

MCM recruitment assay was performed identically to MCM loading assay, with the exception that 5 mM ATP was replaced with 5 mM ATPγS. After the incubation, the beads were washed twice with an ice-cold low salt wash for 20 secs.

### CDK phosphorylation of ORC

For CDK phosphorylated ORC, 12 nM of ORC was incubated with 12 nM CDK for 10 mins at 30°C in loading buffer containing 0.5 mM ATP. To stop the reaction, 60 nM Sic1 was added and incubated for an additional 5 mins at 30°C. For non-phosphorylated ORC, 12 nM ORC was first incubated with 60 nM Sic1 for 5 mins at 30°C and then with 12 nM CDK for 10 mins at 30°C. The subsequent loading assay was done as described above. For Fig. 1b, proteins were separated by Criterion XT Tris Acetate 3-8% (Bio-rad) SDS-PAGE and visualized by silver staining.

### Salt sensitivity assay

The salt sensitivity assay was performed identically to the MCM loading assay. After 20 min incubation, the samples were washed twice with cold LSW containing the indicated NaCl concentration, as shown in Fig. 6a, for 30 sec, followed by two washes with cold LSW.

### Retention assay

Retention assay was performed identically to the MCM loading assay. After 20 min of incubation, the beads were washed with HSW (for 90bp origin only) and then cold loading buffer (for both one site and 90bp origins) for 40 sec at 30°C with mixing at 1600 rpm. 50X molar excess of competitor DNA in loading buffer (preheat at 30°C) was then added and the mixture was aliquoted for an additional wash with HSW (for 90bp origin only), followed by cold LSW (for both one site and 90bp origins) at the indicated times.

### Replication assay

The replication assays were performed according to a protocol described by Yeeles *et al.*, 2015^47^. 40 nM ORC or phosphorylated ORC, 40 nM Cdc6, 60 nM MCM/Cdt1 and 4 nM DNA were incubated for 20 mins at 30°C, 1250 rpm in reaction buffer containing 25 mM HEPES-KOH pH7.6, 10 mM MgOAc, 2 mM DTT, 0.02% NP-40, 100 mM KGlu, 5 mM ATP. The ORC was phosphorylated after CBP pull-down during purification and excess CDK was removed through gel filtration. 50 nM DDK was added with further incubation for 10 mins. The reaction was then incubated with 40 nM Cdc45, 30 nM Dpb11, 20 nM Pol ε, 20 nM GINS, 20 nM CDK, 100 nM RPA, 20 nM Ctf4, 10 nM TopoI, 30 nM Pol α, 25 nM Sld3/7, 20 nM Mcm10, 10 nM Sld2, 33 μM ^32^P-dCTP, 200 μM NTPs and 80 μM dNTPs for 30 mins, 30°C, 1250 rpm. The reactions were then stopped by adding 50 mM EDTA (final concentration) and filtered through an Illustra Microspin G-50 column. The replication products were resuspended in loading buffer and separated through 0.8% alkaline agarose gels in 30 mM NaOH, 2 mM EDTA for 16 hrs at 25 volts. Gels were fixed with 5% cold trichloroacetic acid and then dried onto chromatography paper (Whatman) and autoradiographed with Amerhsam Hyperfilm-MP (GE Healthcare). Gel images were scanned using a Typhoon phosphorimager (GE Healthcare) and were quantified using ImageJ.

### EMSA

Double-stranded DNA probes for electrophoretic mobility shift assays (EMSA) were generated by PCR using primers (50bp up 209 and 50bp down 209) to give 265bp probes. A full list of primers and plasmids used for DNA templates are listed in Supplementary Materials.

62.5ng gel-purified PCR product (Nucleospin Gel and PCR clean-up, 740609.250 Macherey Nagel) was 5’ end labelled in a final volume of 10 μl using 0.6 μl T4 PNK (10U/μl, M0201 New England Biolabs) with 2 μl gamma 32P ATP 3000Ci/mmol (SRP-301 Hartmann). After 1 h on ice, 40 μl 10 mM HEPES-KOH pH7.6, 1 mm EDTA was added and unincorporated nucleotides were removed by passing through 2x microspin G-50 columns (GE27-5330-02 Cytiva), pre-spun for 1’ at 735g followed by elution for 2’ at 735g.

For each EMSA reaction, 12 fmols DNA probe was used (assuming 80% elution losses per column). Reaction components were added on ice to give a final concentration of 25 mM HEPES-KOH pH7.6, 10 mM MgOAc, 100 mM NaOAc, 0.02% NP40, 5 mM DTT, 80 mM KCl, 5% glycerol, 2 μg/ml poly dI-dC and 5 mM ATP. Lastly, 100 fmol of purified ORC was added as required and the reactions were incubated for 30 mins at 30°C in a thermomixer at 800 rpm. After 30 mins the samples were placed on ice and 2 μl loading dye (Purple gel loading dye, no SDS, B7025 New England Biolabs) was added prior to loading 12 μl onto a 1.5% 15 x 15 cm 100 ml 0.5X TB agarose gel. The gel was run for 1h at 200 V (4°C) before fixation for 30 mins in 5% TCA and drying on a vacuum gel dryer at 55°C for 1h (Hoefer GD2000). After overnight exposure to a storage phosphor screen (BAS-IP MS 2025 E GE Healthcare) the signal was detected using a Typhoon FLA7000 phosphoimager.

### Induction of re-replication

Cells were grown overnight in medium lacking methionine and containing 2% raffinose (1X YNB, 1X CSM-Met (Formedium, DCS0111). At a density of about 1×10^7^ cells/ml, 50 ng/ml alpha factor was added to arrest the cell growth for 3 hrs. Cells were spun down and switched to YPRaff medium with alpha factor for 1 hr. Cells were washed 2X with YPRaff media (1X with YPRaff and 1X with YPRaff + 50 μg/ml Pronase E (Sigma-Aldrich, 1074330001)). Cells were resuspended in YPRaff medium containing 100 μg/ml Pronase E with 5 μg/ml nocodazole for 3 hours. Cells were then divided into two and 2% of glucose or galactose was added, respectively, to induce Cdc6ΔNT for 3 hrs in the presence of galactose only. Cells were collected and lysed with lysis buffer (2% Triton, 1% SDS, 100 mM NaCl, 10 mM Tris, pH8.0, 1 mM EDTA and protease inhibitor (AEBSF, Leupeptin, Pepstatin A). Glass beads and phenol/chloroform were added together and tubes were vortexed for 1 min. The supernatant was treated with RNases at RT for 30 min and then extracted with phenol/chloroform and ethanol precipitated. The DNA was used in qPCR. 9 μl of qPCR reaction was set up in 384 well plates with 4.5 μl of 2X FastStart Universal SYBR Green Master mix ROX (Roche), ∼0.28 μM of primers (Set 1: CTL141+CTL142 for re-replication region; Set 2: CTL143+CTL144 for internal control) for 10 min at 95°C and then 40 cycles of 10s at 95°C and 30s at 60°C. Each DNA sample was first evaluated for the linearity of PCR by performing serial dilutions. The fold change of threshold cycle (Ct) was determined using 2^^-ΔΔCt^ method^48^, where ΔΔCt = (Ct_re-replication region_ – Ct_internal control_)_galactose_ – (Ct_re-replication region_ – Ct_internal control_)_glucose_

### Nucleosome flanked origins

Nucleosome-flanked origins of replication were prepared with purified yeast histone octamers as described^11^ on the DNA substrates of N-ARS1-N or N-90bp-N. Plasmids containing N-ARS1-N and N-90bp-N were synthesised by Eurofins and used as a template for PCR to produce linear DNA substrates using the primers NCP F and NCP R. Origin substrates were amplified using standard PCR protocols and were purified by anion exchange chromatography using a 1 ml RESOURCE Q column (GE healthcare). Peak fractions were ethanol precipitated and resuspended in TE buffer.

### Nucleosome assembly

Purified origin DNA substrates were combined with soluble histone octamers to form nucleosomes by salt-deposition ^11,49^. Nucleosome reconstitution was optimized in small-scale titrations and the products checked by 4% native PAGE.

### Negative stain EM MCM loading assays

MCM loading experiments, negative stain sample preparation, imaging and analysis were carried out as described^11^, with minor modifications:

### Sample preparation

Nucleosome flanked ARS1/90bp origin substrates (7.5 nM) were incubated with wild-type or mutant ORC (20 nM), Cdc6 (20 nM) and MCM-Cdt1 (40 nM) with mixing (1250 RPM) in EM buffer (25 mM HEPES pH 7.6, 10 mM MgOAc, 100 mM NaOAc, 0.02% NP-40, 80 mM KCl, 1 mM DTT and 5% glycerol) and 2 mM ATP in a final reaction volume of 20 μl. In experiments testing CDK-phosphorylated ORC, ORC was pre-phosphorylated for 10 minutes before Sic1 was added for 5 minutes to inhibit further CDK activity. Negative stain grids were prepared after 2-minutes (MO detection) and 10-minutes (DH detection) incubation at 24°C. In all cases, samples were diluted 5-fold prior to making grids.

### Grid preparation

Samples were applied to glow-discharged 300-mesh copper grids with carbon film (EM Resolutions). 3 μl of sample was applied to each grid and incubated for 1 minute. Staining was performed with four drops of 40 μl 2% uranyl acetate, before grids were blotted to remove excess stain.

### Data collection

Micrographs were collected on a Tecnai LaB6 G^2^ Spirit transmission electron microscope (FEI) operating at 120 keV, with a 2K x 2K GATAN Ultrascan 100 camera. Images were recorded at a nominal magnification of 30K (3.45 Å pixel size), and a defocus range of ∼0.5 – 2 μm.

### Image processing

All datasets were processed in Relion3.1^50^ using a common processing pipeline. The contrast transfer function (CTF) of each micrograph was estimated using Gctf v1.06^51^. Particles were picked with Topaz^52^ using a model that had been pre-trained to pick MCM-Cdt1, ORCs, MOs and MCM-DHs in negative stain EM data. Particles were initially extracted with a 128-pixel box, rescaled to 64 pixels and were subjected to two rounds of reference-free 2D classification. All MCM containing particles were re-extracted with a 128-pixel box without rescaling. The particles were subjected to a further round of 2D classification. From this classification, particles that contributed to classes containing the MO and DH intermediates were selected and further classified, as required. Finally, particles contributing to classes that contained an MCM (total MCMs, MOs and DHs) were quantified from the appropriate 2D classification. The proportion of MCM-containing particles contributing to each intermediate state was plotted using Prism.

### ReconSil

Datasets for ReconSil images were processed together in Relion3.1^50^. CTF estimation was performed using Gctf v1.06^51^. Particles were picked with Topaz^52^ using models pre-trained to pick MCM-containing particles (as above), ORCs and nucleosomes. Particle picks were combined, duplicates removed and extracted particles were subjected to multiple rounds of reference-free 2D classification. ReconSil micrographs were generated using the command-line tool ‘relion_particle_reposition’ in Relion3.1 to overlay particles in the raw micrographs with the 2D averages that those particles contributed to. Individual origins were extracted from raw and ReconSil micrographs (box size: 320 pixels). Representative examples were selected from fully reconstituted origins where confident assignment of co-localisation to the same origin could be made (as described in^11^).

### Cryo-EM

Image acquisition, refinement and validation statistics for the experiments below can be found in **Extended Data Table 1**.

### Sample preparation

Samples were prepared as described above for the negative stain EM experiments with the following modifications:

Mutant ORC (Orc6-4A; 40 nM) was phosphorylated by CDK (10 nM) for 10 minutes in CDK buffer (25 mM HEPES pH 7.6, 10 mM MgOAc, 100 mM NaOAc, 80 mM KCl, 1 mM DTT, 2 mM ATP), before Sic1 (30 nM) was added to inhibit further CDK activity. Nucleosome flanked ARS1 origin substrates (15 nM) were incubated with pre-phosphorylated Orc6-4A mix (as above) and MCM-Cdt1 (80 nM), with mixing (1250 RPM) in cryo-EM buffer (25 mM HEPES pH 7.6, 10 mM MgOAc, 100 mM NaOAc, 80 mM KCl, 1 mM DTT) and 2.2 mM ATP in a final reaction volume of 50 μl. The reaction was incubated at 30°C for 30 minutes before being used to prepare cryo-EM grids. 4 μl of sample was applied to fresh graphene-oxide coated 300-mesh UltrAuFoil R1.2/1.3 grids^53^ and incubated for 30 seconds before vitrification using a Vitrobot Mark IV (Thermo Fisher) cooled to 10°C with 100% humidity. Grids were blotted for 5 seconds and plunged into liquid ethane.

### Cryo-EM data collection

Data were collected on a Titan Krios EM equipped with a K2 Summit direct electron

detector (Gatan Inc.) at the Francis Crick Institute (Structural Biology STP).

### Cryo-EM image processing: SH and DH

Image processing was performed in RELION-3.1^50^ (Extended Data Figs. 7 & 8). Movie stacks were aligned and motion-corrected using MotionCor2^54^, retaining all frames. The contrast transfer function (CTF) of each micrograph was estimated using Gctf v1.06^51^. Particles were automatically picked using Topaz^52^. First, particles were picked using a general Topaz model (scale factor = 4; model = resnet8_u32; radius = 20). These particles were extracted (320-pixel box, rescaled to 80 pixels) and subjected to multiple rounds of reference-free 2D classification to isolate SH and DH particles for subsequent processing.

High-quality DH particles were subjected to iterative rounds of 3D refinement, particle polishing (incl. increasing the box size to 420 pixels), CTF refinement, and a final round of 2D classification, yielding the final EM map at an average resolution of 2.8 Å, reconstructed from 135,143 MCM-DH particles. The final RELION map was subject to density modification using resolve_cryo_em in Phenix 1.19.2^55^ to generate a 2.6 Å map used for model building (Extended Data Fig. 5).

SH particles obtained from 2D classification were used for Topaz training (scale factor = 4; cnn_model = resnet8; radius = 3). SH particles were picked using the new Topaz model (selection threshold = -2), extracted (280-pixel box, rescaled to 70 pixels) and subjected to two-rounds of reference-free 2D classification. SH particles were re-extracted (280-pixel box, unbinned) and classified into 3 classes using a 30-Å filtered reference from an ab-initio reconstruction in CryoSPARC^56^. High-quality SH particles were subjected to iterative rounds of particle polishing (incl. increasing the box size to 420 pixels), CTF refinement, masked 3D refinement, and a final round of 2D classification (without alignment), yielding the final EM map at an average resolution of 3.4 Å, reconstructed from 396,366 SH particles. The final RELION map was subject to density modification using resolve_cryo_em in Phenix 1.19.2^55^ to generate a 3.1 Å map used for model building (Extended Data Fig. 5).

### Molecular modelling

#### DH

Molecular modelling of the DH complex was performed using a density modified map generated using resolve_cryo_em. A published DH model (PDB 7P30^57^) was used as an initial model and rigid-body docked into the new DH EM map. The resulting model was subject to manual modification in Coot^58^ and real space refinement in Phenix 1.19.2^59^. All figures were generated using UCSF ChimeraX^60^.

#### SH

Molecular modelling of the SH complex was performed using a density modified map generated using resolve_cryo_em. An initial model of a loaded SH was extracted from the higher-resolution DH model (this study) and rigid-body docked into the SH EM map. The resulting model was subject to iterative rounds of manual modification in Coot^58^ and real space refinement in Phenix 1.19.2^59^. All figures were generated using UCSF ChimeraX^60^.

### Orc2 IDR AlphaFold2 Modelling

The MCM-ORC (MO) interaction interface, consisting of *S. cerevisiae* Orc2 (P32833), Orc3 (p54790), Orc5 (P50874, aa 319-479), Orc6 (P38826), Mcm2 (P29469, aa 178-457), Mcm5 (P29496 aa 1-336) and Mcm6 (P53091, aa 88-463), was modelled using AlphaFold2 (2.3.1^13^) implemented in AlphaPulldown (0.30.0^61^). The five resulting models positioned the Orc2 IDR amino acids 190-231 in the same location, wrapping around the Orc6 TFIIB-B domain. Orc3 (aa 270-419) in the top ranked model was aligned with the corresponding sequence in the previously determined MO structure (6RQC^11^) using the matchmaker tool in ChimeraX (v1.6^59^). The Orc2 IDR (aa 190-231) was fit as a rigid body into the previously unassigned density in the MO map using the ‘Fit in Map’ tool in ChimeraX. Fig. 1c and Extended Data Fig. 1 (AlphaFold2 figure) display EMready^62^ density modified EM maps obtained from the multibody refinement of the MO intermediate. Similar AlphaFold2 predictions of the ORC subunits alone, or the ORC subunits with the complete N-terminal domains of Mcm2-7, yielded almost identical predictions for the interaction of the Orc2 IDR with Orc6.

### Simulating DH formation efficiency from two diffusing SHs

We model the separation dependent DH formation efficiency using stochastic simulations of 1-dimensional diffusion of SHs sliding along the DNA. DH formation efficiency is computed from the probability of collision of two diffusing SHs given by intersection of their diffusion trajectories. The simulation consists of four key steps: SH loading, SH diffusion, SH dissociation, and DH formation,

#### 1. SH loading

The simulation takes place on a DNA substrate consisting of two ORC binding sequences in a head-to-head orientation with the B1 sequences of the EACS/B1 (ORC binding site) pointed towards each other. The separation between the two origins is defined as the distance between end of the B1 sequences in ORC binding sequences as shown in Extended Data Fig. 9a. ORC binds to an EACS/B1 sequence with an overhang of 6 base pairs beyond the B1^14,63^ (Extended Data Fig. 9b). This ORC complex can then load an SH ahead of itself which occupies 35 base pairs on the DNA substrate^64^. This implies that the loading of one SH complex onto the DNA occupies at least 41 base pairs (6 bp ORC overhang + 35 bp SH) beyond the origin on the DNA substrate. The simulation is initialized with both ORC sites loaded with SHs with N-termini pointing towards each other.

#### 2. SH diffusion

Since SH sliding is independent of ATP hydrolysis, we assume that its diffusion occurs at thermal equilibrium and is therefore not driven by input of external energy. In such conditions, the displacement of SH per unit time follows a gaussian distribution given by:

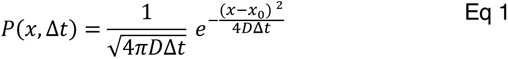

where, *x* is the position of the SH, *D* is the diffusion coefficient, Δ*t* is the duration of the time step and *x*_0_ is the initial position. After initialization of the simulation, the displacement of SH in the following time step is determined by sampling this probability distribution using the value of SH diffusion constants measured from single molecule fluorescence tracking experiments_31,32_.

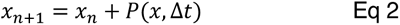

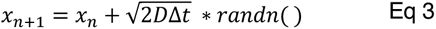

where ‘randn()’ is the MATLAB function to generate normally distributed random numbers. The time step of simulation is chosen such that the mean squared displacement (MSD) of SH for the given value of diffusion constant is less than 1 base pair of the DNA.

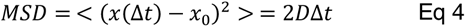

Thus, the time step of the simulation was chosen to be 10 ms.

#### 3. SH dissociation from the DNA

Once loaded onto the DNA substrate, SHs dissociate form the DNA with an experimentally measured half-life of 3 minutes (Fig. 5b). We implement this phenomenon in our simulations by assigning a dissociation probability to each loaded SH onto the DNA. The dissociation probability is computed using the exponential distribution that satisfies the experimentally measured half-life:

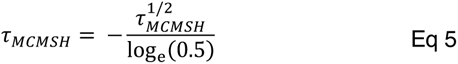

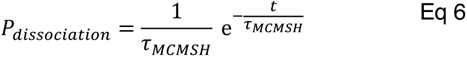

where 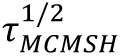 is the experimentally measured SH half-life and *t* is the total time spent by SH on the DNA since its loading. At every time step, a random number, R is sampled from a uniform distribution using MATLAB function rand(). If R is smaller that *P*_*dissociation*5677896:)68+_, the corresponding SH is said to be dissociated from the DNA, and the simulation is stopped as DH formation is no longer possible.

#### 4. DH formation

To successfully form a DH, the two loaded SHs must collide with each other while diffusing along the DNA. We define a collision event as the first instance when the distance between the N-termini of both SHs is less than or equal to zero base pairs during the simulation. The time duration from the start of the simulation to the time step corresponding to the collision event is referred to as the collision time which also corresponds to the first-passage time of the collision process. The probability of DH formation is thus the probability of collision of two SHs loaded onto the DNA which depends on the separation between the ORC binding sites, the availability of binding site onto the DNA for the second SH, the diffusion constants of the SHs, the half-life of SH onto the DNA and the total time for the simulation.

### Simulation Results

The diffusion coefficient of MCM onto the DNA has been observed to be 800 ± 200 *bp*^2^/*s* from single-molecule fluorescence tracking experiments^32^. The half-life of SH on the DNA is measured to be 3 minutes (Fig. 5b). Using these values as our simulation parameters, we quantify the dynamics of DH formation as a function of the separation between the ORC loading sites as follows. For each case of separation, we first quantify the number of simulated trajectories that contain a collision event between two SHs and measure the time between the start of the simulation to the collision event, hereafter referred to as collision time (*t_c_*) Extended Data Fig. 10a. The probability distribution of collision times, *P*(*t_c_*), is generated from all the iterations (Extended Data Fig. 10b). The cumulative sum of this distribution given by 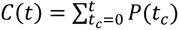, refers to the probability of collisions occurring before time *t*, which is by definition, the probability of DH formation within time *t* (Extended Data Fig. 10c). Computing *C*(*t*) for various separations between ORC binding sites at a fixed time (20 minutes) determined by the incubation time in bulk experiments (Fig. 5d) allows us to compare the relative DH formation efficiencies on the corresponding DNA substrates (Extended Data Fig. 10d).

We observe that the relative DH formation efficiency has a maximum at separation of 82 base pairs between the ORC binding sites, which corresponds to the case when SHs are loaded closest to each other onto the DNA (ORC overhang + SH + SH + ORC overhang = 6+35+35+6 = 82 bp). At separations greater than 82 bp, the DH formation efficiency decreases with increasing separation (and shows a good agreement with exponential trend) as both SHs loaded onto the DNA must diffuse along the DNA and collide with each other before either or both dissociate from the DNA.

## Acknowledgements

We thank A. Alidoust, N. Patel, and D. Patel in the Structural Biology STP for yeast protein expression. We thank A. Nans, A. Purkiss and P. Walker from the Structural Biology STP for support with cryo-EM and computing. We thank G. Cameron and H. Yardimci for the gift of Lac repressor protein and plasmid. We thank F. Palmero Moya of the NHD lab for discussions of simulations. This work was supported by the Francis Crick Institute, which receives its core funding from Cancer Research UK (FC001065 and FC001066), the UK Medical Research Council (FC001065 and FC001066), and the Wellcome Trust (FC001065 and FC001066). This work was also funded by Wellcome Trust Senior Investigator Awards (106252/Z/14/Z and 219527/Z/19/Z) to JFXD, and European Research Council Advanced Grants (669424-CHROMOREP and 101020432-MeChroRep) to JFXD. AC receives funding from the European Research Council (ERC) under the European Union’s Horizon 2020 research and innovation programme (grant agreement no. 820102). NHD acknowledges funding from the Netherlands Organization for Scientific Research (NWO) through grant OCENW.M.21.173 and from a European Research Council Advanced Grant (789267-REPLICHROMA). T.C.R.M. was supported by a Novo Nordisk Fonden Hallas-Møller Emerging Investigator Grant (NNF22OC0073571), the Danish National Research Foundation (DNRF115) and the Carlsberg Foundation (CF21-0571). For the purpose of Open Access, the author has applied a CC BY public copyright licence to any Author Accepted Manuscript version arising from this submission.

## Author contributions

CTL and JFXD conceived the initial study. CTL performed all biochemical MCM loading experiments, analysis of re-replication *in vivo* and helped with EM experiments. TCRM performed all EM experiments, image processing and atomic model building. AC helped with AlphaFold-Multimer, cryo-EM grid preparation and screening. KWT performed *in vitro* replication experiments. AE performed the EMSA experiment. PE performed the PWM analysis of yeast origins. ST, HS, CTL, JFXD and NHD conceived the simulation experiment which was developed and executed by ST with advice from HS and NHD. CTL, TM and JFXD wrote the manuscript with input from other authors.

## Data availability

Atomic model coordinates and cryo-EM maps have been deposited in the Protein Data Bank (PDB) and Electron Microscopy Data Bank (EMDB), under the accession codes 8RIF/EMD-19186 (DH), and 8RIG/EMD-19187 (SH).

## Extended Data

**Extended Data Figure 1.**
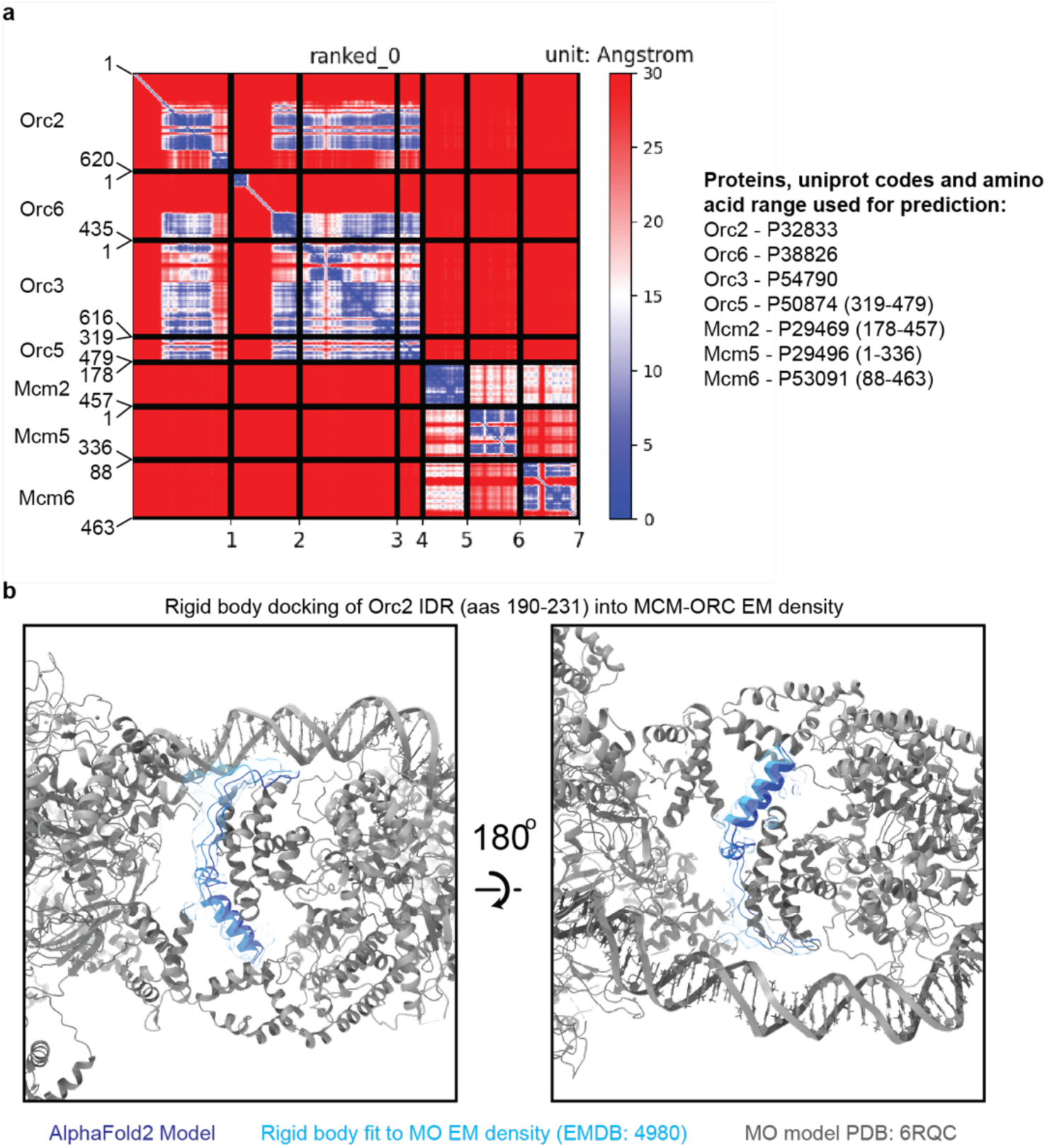
AlphaFold2 places the Orc2 IDR at the MCM-ORC (MO) interface. **a.** Predicted aligned error (PAE) plot for the AlphaFold2 model of the MO interface. The structure prediction was based on proteins present at the MO interface in the published structure (PDB: 6RQC^11^). **b.** Following alignment of Orc3 (aa 270-419) in the top ranked AlphaFold2 model with the respective region of Orc3 in the MO structure, the predicted Orc2 IDR (aas 190-231) was fit as a rigid body into previously unassigned EM density in the MO map. Here, the density displayed has been isolated from a density modified (EMready^62^) map of ORC obtained from the multibody refinement of the MO (EMDB: 4980^11^). The prediction places the Orc2 IDR as wrapping around the Orc6 TFIIB-B domain at the center of the MO interface.

**Extended Data Figure 2.**
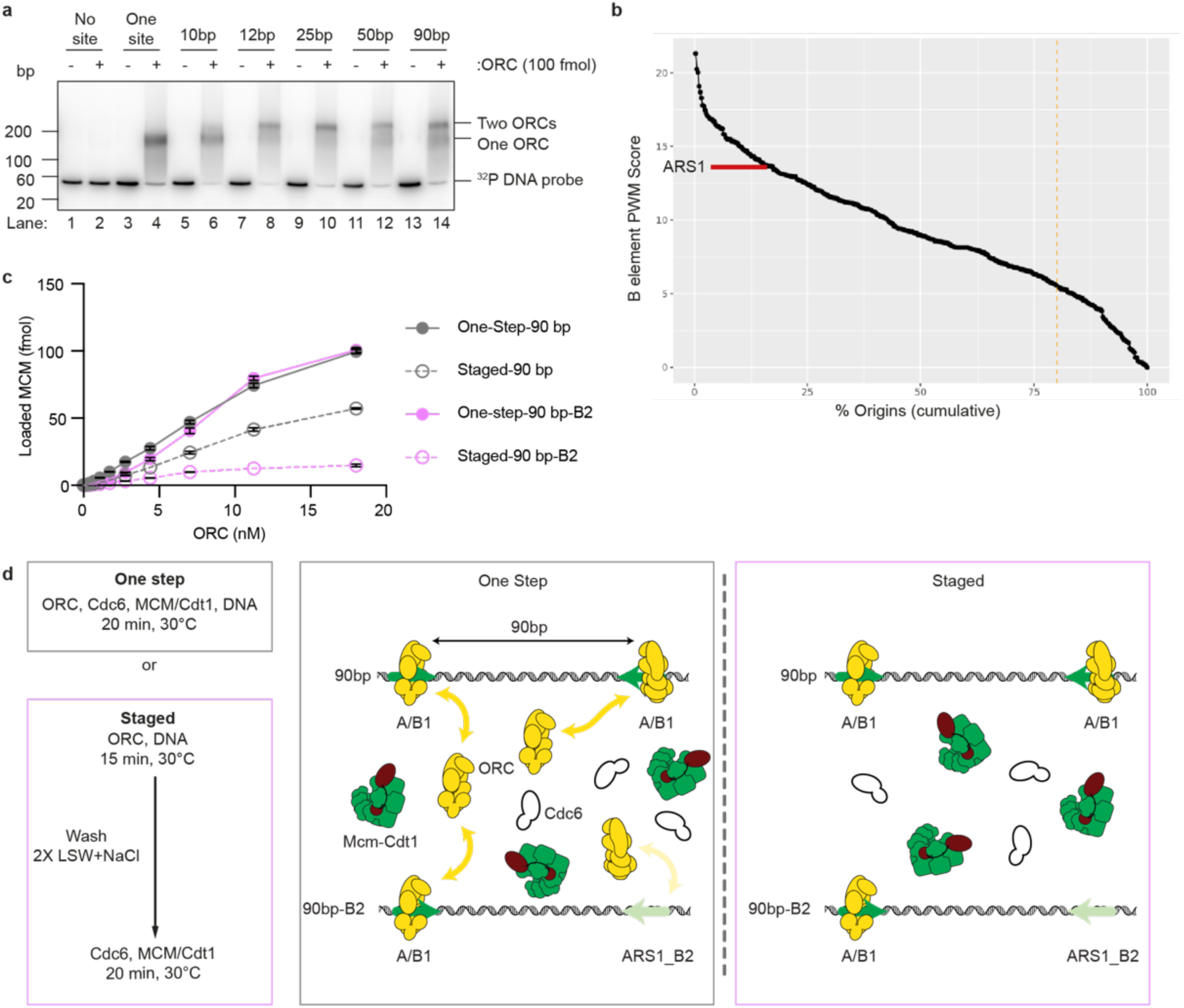
The formation of MO stabilizes ORC binding at weak binding sites. **a.** EMSA of ORC binding at different distances on synthetic templates. The 10bp distance is similar to ARS1, which can only bind a single ORC. Two ORCs can bind simultaneously on origins with a spacing of ≥12bp. b. Approximately 15% of origins have a better match to the EACS at the weaker ORC binding site than ARS1 B2. c. Near-saturating amounts of MCM are loaded onto both 90bp and 90bp-B2 origins in one-step reactions using 18 nM of ORC. d. Schematic figure of ‘One Step’ and ‘Staged’ reactions, where excess ORC is always available ifor the one step reaction, but ORC only remains bound to high affinity sites after stringent washing with 80 mM NaCl in the staged reactions.

**Extended Data Figure 3.**
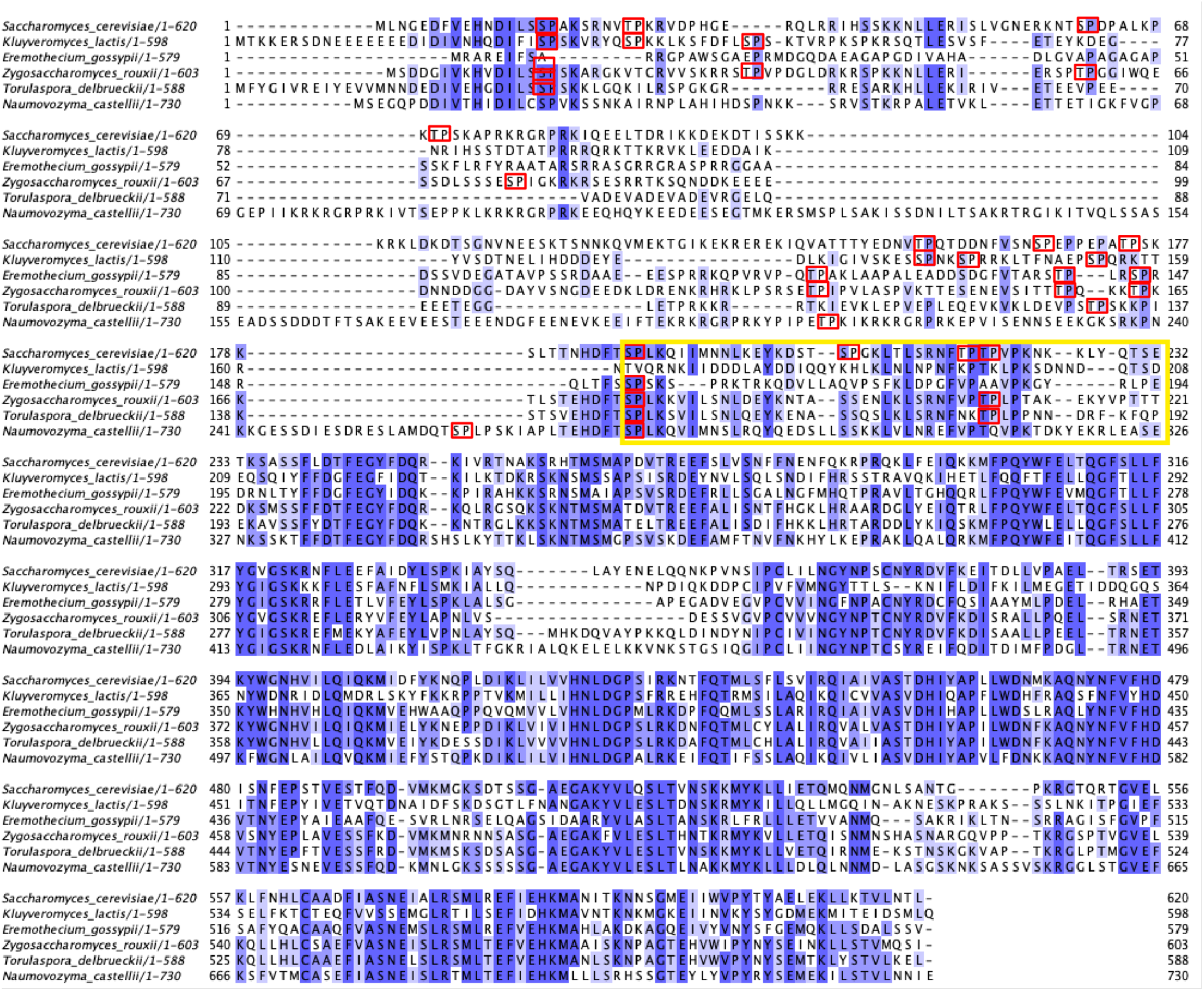
Closely related yeast species contain multiple potential CDK phosphorylation sites in the Orc2 IDR. The region of the Orc2 IDR that is predicted to form part of the MCM-ORC interface (yellow box) is moderately conserved and harbors multiple potential CDK phosphorylation sites.

**Extended Data Figure 4.**
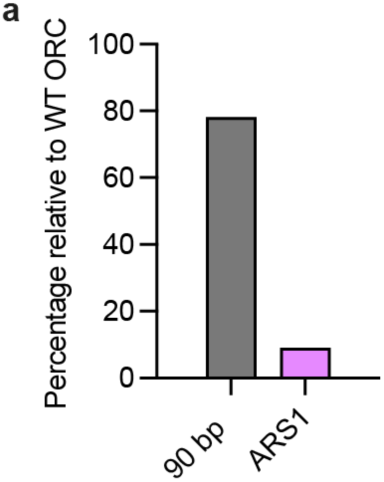
CDK phosphorylated ORC is replication competent in vitro. **a.** The percentage of DNA synthesis for CDK-phosphorylated ORC relative to WT ORC as shown in Fig. 3c.

**Extended Data Figure 5.**
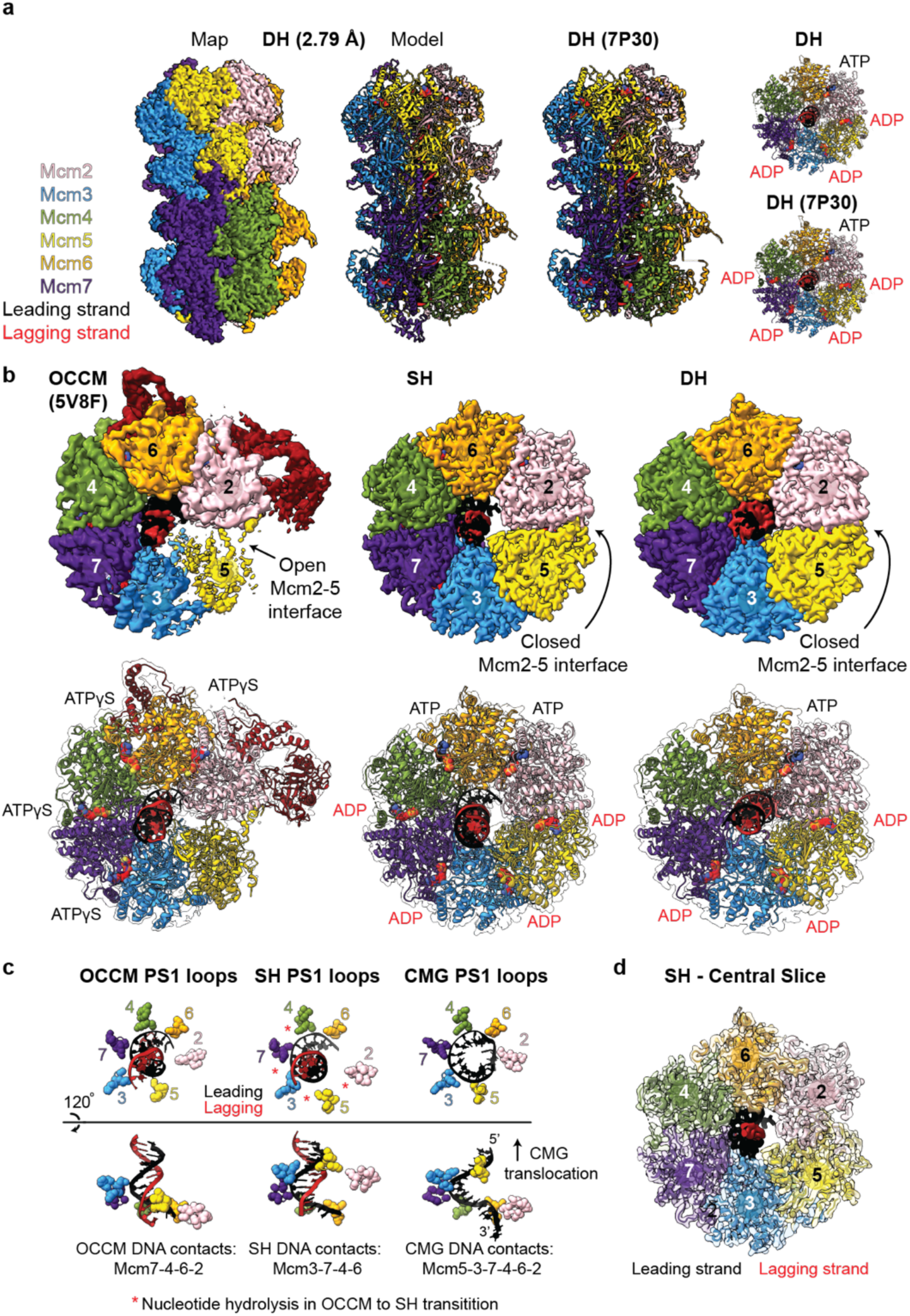
Comparison of DHs assembled by ORC and ORC phosphorylated on Orc2. **a.** Cryo-EM structure and atomic model for DHs assembled by ORC phosphorylated on Orc2 (left; this study) and atomic model for DHs assembled by unphosphorylated ORC (right; PDB: 7P30^57^). The DH EM density map displayed has been density modified using EMReady^62^. **b**. Comparison of the MCM ring in the OCCM (left), SH (middle) and DH (right), showing that the SH and DH have fully closed Mcm2-5 interfaces and both appear to be in post-ATP hydrolysis states relative to the OCCM. **c.** Comparison of the PS1 loops of the OCCM (5V8F^65^; left), SH (this study; middle) and CMG (6SKO^66^; right), showing that the SH PS1 loops adopt a staircase configuration and engage the backbone of the leading strand DNA template like the CMG. **d.** Central slice through the SH structure, highlighting the well resolved EM density for the bases in the leading strand DNA template.

**Extended Data Figure 6.**
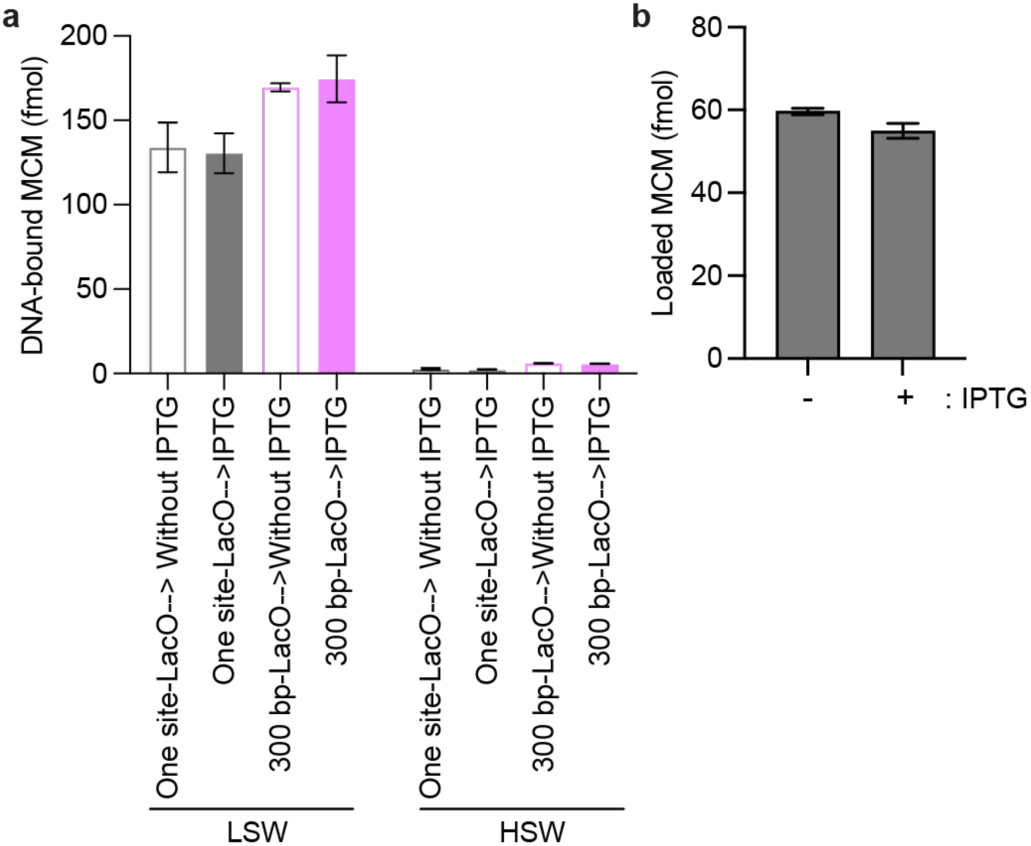
DH formation by two single hexamers. **a.** MCM loading assay performed with 20 minutes incubation of Orc2ΔN, Cdc6 and MCM-Cdt1 with ATP and a DNA template containing two high affinity ORC sites separated by 300bp and 4 lac operators. Aliquots taken before IPTG addition show MCM loaded onto DNA (detectable after LSW), but no HSW stable DH formation. **b.** The addition of IPTG in the MCM loading assay does not affect MCM loading.

**Extended Data Figure 7.**
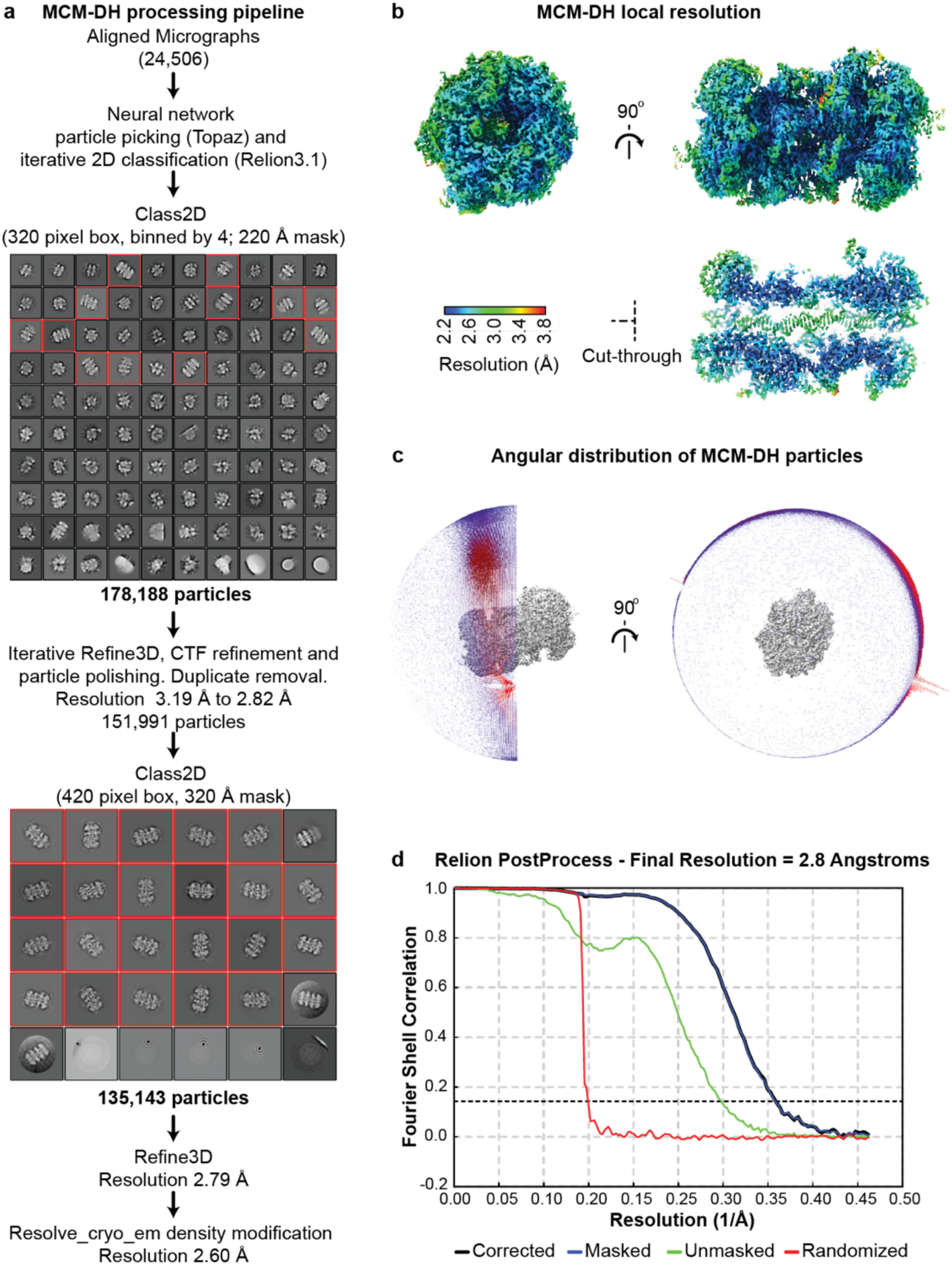
DH cryo-EM data processing. **a.** Cryo-EM data processing pipeline for the DH. b. Final DH EM density map following density modification with resolve_cryo_em^55^, coloured according to the local resolution. c. Angular distribution plot. d. Fourier shell correlation (FSC) plot obtained from Relion PostProcess.

**Extended Data Figure 8.**
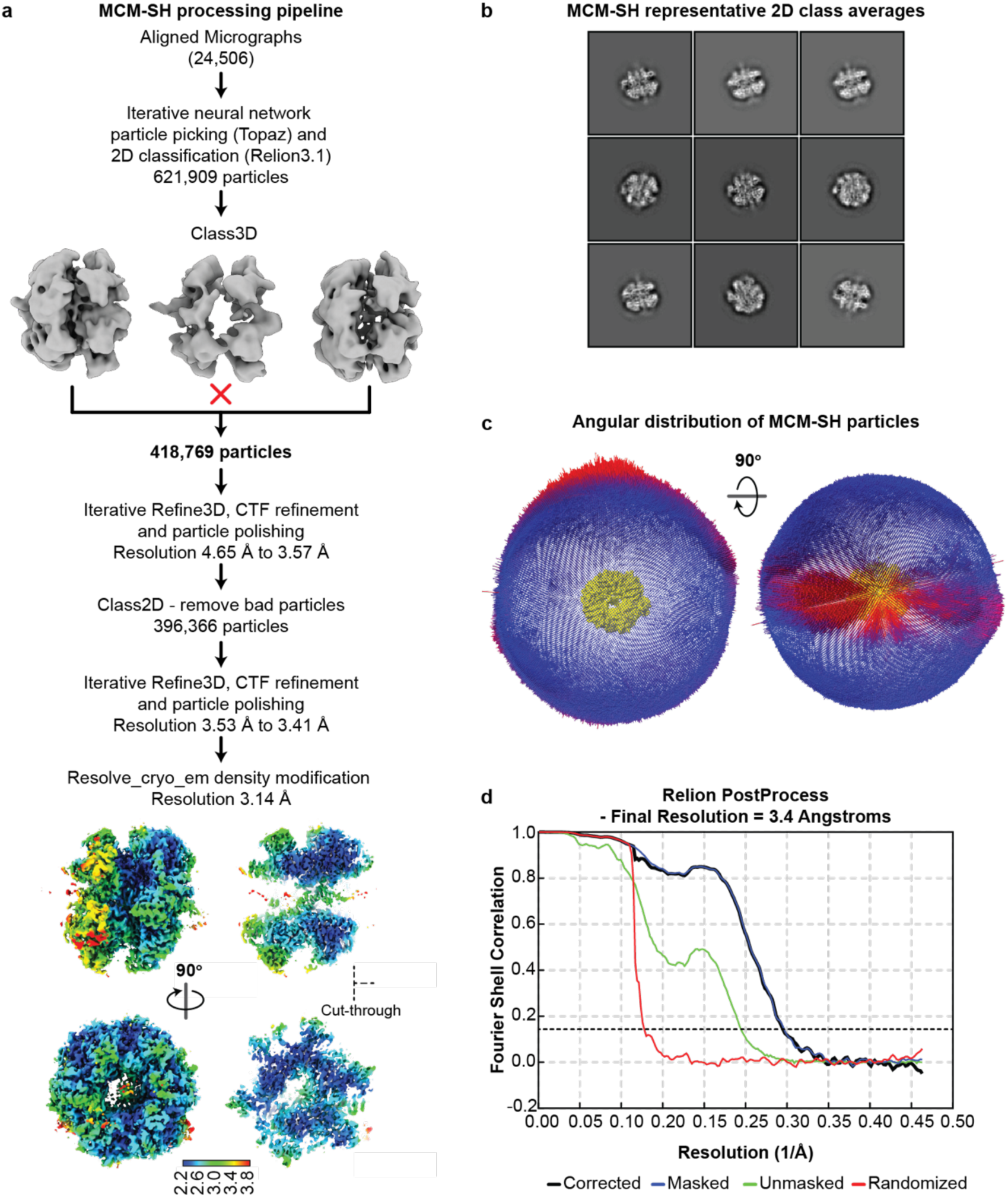
SH data cryo-EM processing. **a.** Cryo-EM data processing pipeline for the SH, including views of the resulting 3D density modified map coloured according to local resolution. b. Representative 2D classes for the SH. c. Angular distribution plot. d. Fourier shell correlation (FSC) plot obtained from Relion PostProcess.

**Extended Data Figure 9.**
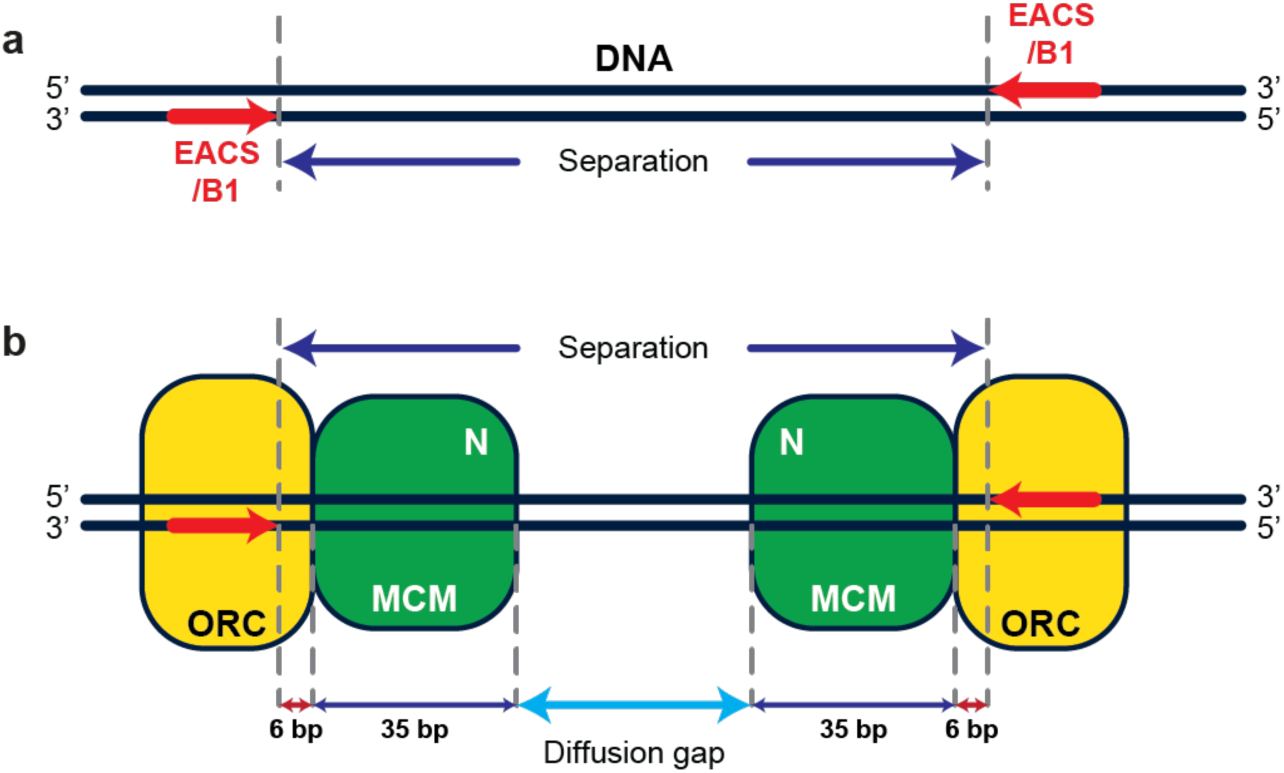
Schematic of the simulated system and definition of relevant distances. **a.** The separation between two ORC binding sites oriented towards each other on DNA is defined as the distance in base pairs between the end of the B1 sequences in the EACS/B1 (‘ORC binding sites’). b. ORC binds to the EACS/B1 sequences with an overhang of 6 bp beyond the end of the B1 sequence. A SH binds in front of the ORC and occupies 35 bp with the distance between the origin and the N-termini of SH being 6+35 = 41 bp. The distance between the N-termini of SHs is the diffusion gap that must be overcome to result in DH formation and is determined by the separation between origin sequences and the space occupied by ORC and MCMs on the DNA.

**Extended Data Figure 10.**
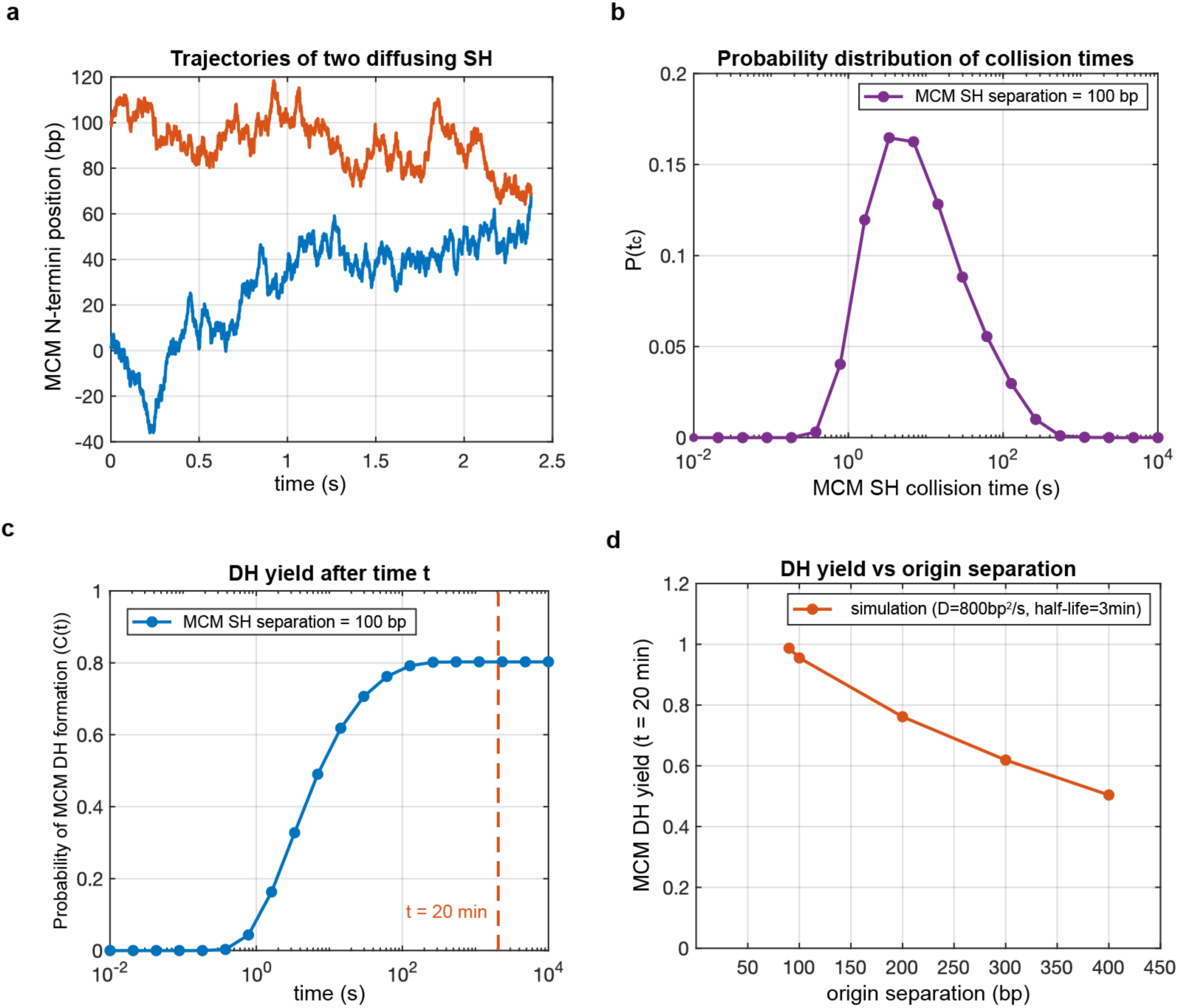
Simulating efficiency of DH formation. **a.** an example of a simulated traces showing two MCM-SHs loaded at a separation of 100 base pairs which then diffuse along the DNA substrate and ultimately collide with each other resulting in MCM-DH formation. b. Probability distribution of MCM-SH collision times. c. Probability of MCM-DH formation with respect to time which is the. Cumulative sum of the MCM-SH collision time distribution shown in (b). MCM-DH yield is defined as the MCM-DH formation probability at a cutoff time determined by the incubation time in vitro experiments (in this case t_cutoff = 20 min). d. Simulated MCM-DH yield at various values of separation between origin sequences.

**Extended Data Movie 1. Model for DH formation via rotation-coupled diffusion of SHs**

Two MCM single hexamers diffusing via rotation along B-form DNA. By using the first steric clash between hexamers as the end point of rotational diffusion, an inter-ring register is obtained which appears close to the register observed in the MCM double hexamer.

**Extended Data Table 1.**

|  | **MCM-SH** (EMDB-19187)<br>(PDB-8RIG) | **MCM-DH** (EMDB-19186)<br>(PDB-8RIF) |
| --- | --- | --- |
| **Data collection and processing** |  |  |
| Magnification | 130,000 | 130,000 |
| Voltage (kV) | 300 | 300 |
| Electron exposure (e-/Å<sup>2</sup>) | 51.4 | 51.4 |
| Defocus range (μm) | 1.0 – 5.0 | 1.0 – 5.0 |
| Pixel size (Å) | 1.08 | 1.08 |
| Symmetry imposed | C1 | C2 |
| Initial particle images (no.) | 621,909 | 178,188 |
| Final particle images (no.) | 396,366 | 135,143 |
| Map resolution (Å) |  |  |
| RELION (PostProcess) | 3.41 | 2.8 |
| FSC threshold | 0.143 | 0.143 |
| Resolve Cryo-EM | 3.14 | 2.60 |
| FSCref threshold | 0.5 | 0.5 |
| Map resolution range (Å) | 2.2-4.0 | 2.2-3.8 |
| **Refinement** |  |  |
| Initial model used (PDB code) | 8RIF | 7P30 |
| Map resolution (Å) | 2.6 | 3.0 |
| FSCref threshold | 0.5 | 0.5 |
| Map sharpening *B* factor (Å<sup>2</sup>) | Resolve Cryo-EM (27.3) | Resolve Cryo-EM (0) |
| Model composition |  |  |
| Non-hydrogen atoms | 30,221 | 64,145 |
| Protein residues | 3,721 | 7,820 |
| Nucleotides | 38 | 106 |
| Ligands | 2 ATP, 4 ADP, | 2 ATP, 6 ADP, |
| (ATP, ADP, Mg<sup>2+</sup>, Zn<sup>2+</sup>) | 6 Mg<sup>2+</sup>, 5 Zn<sup>2+</sup> | 8 Mg<sup>2+</sup>, 10 Zn<sup>2+</sup> |
| *B* factors (Å<sup>2</sup>) |  |  |
| Protein | 65.37 | 61.85 |
| Nucleotide | 20.00 | 102.70 |
| Ligand | 57.35 | 63.70 |
| R.m.s. deviations |  |  |
| Bond lengths (Å<sup>2</sup>) | 0.007 | 0.007 |
| Bond angles (°) | 1.117 | 1.232 |
| Validation |  |  |
| Molprobity score | 1.26 | 1.30 |
| Clashscore | 2.04 | 1.81 |
| Poor rotamers (%) | 0.97 | 0.98 |
| Ramachandran plot |  |  |
| Favored (%) | 95.88 | 94.87 |
| Allowed (%) | 3.96 | 4.98 |
| Disallowed (%) | 0.16 | 0.16 |

## Supplementary Materials

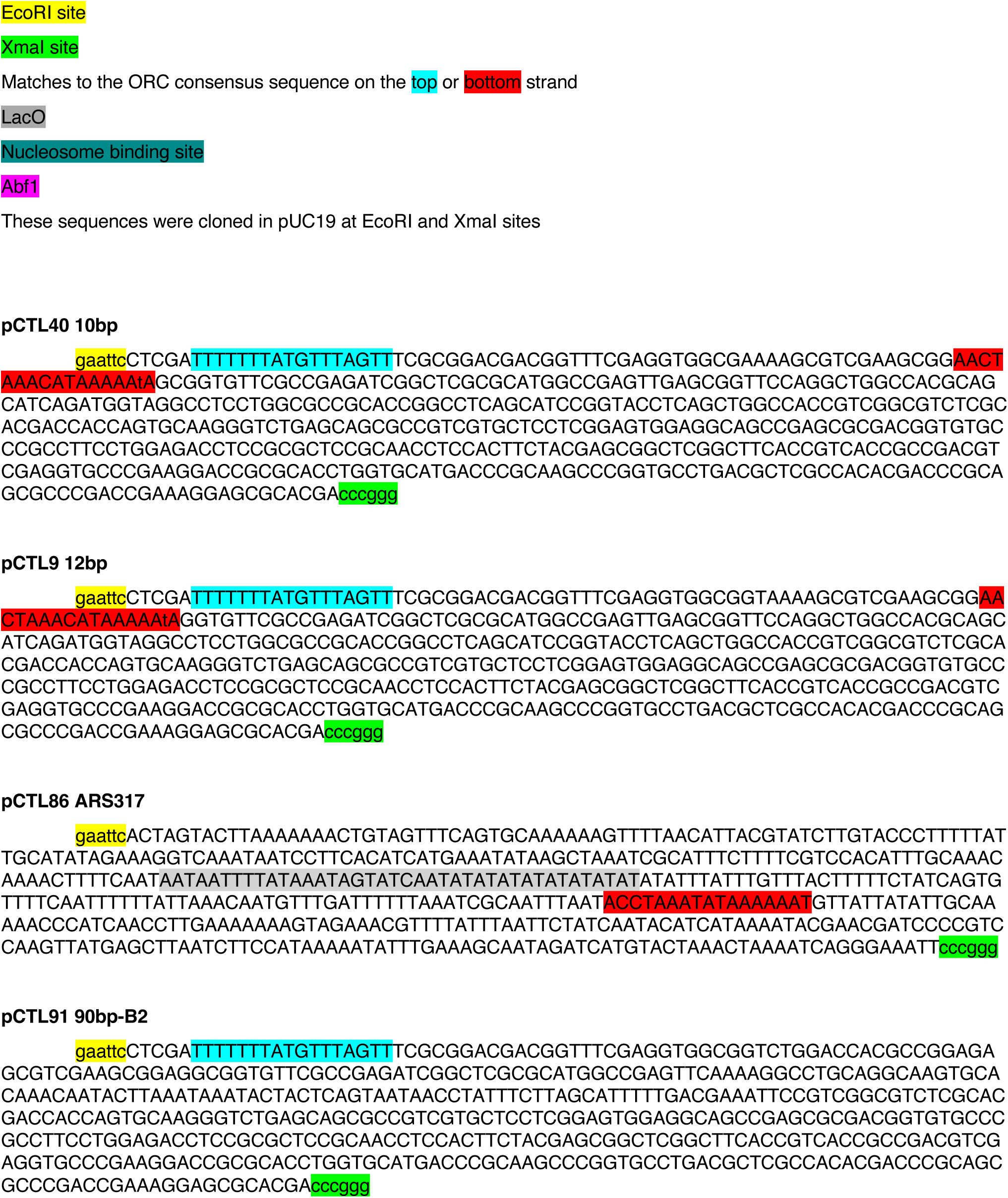
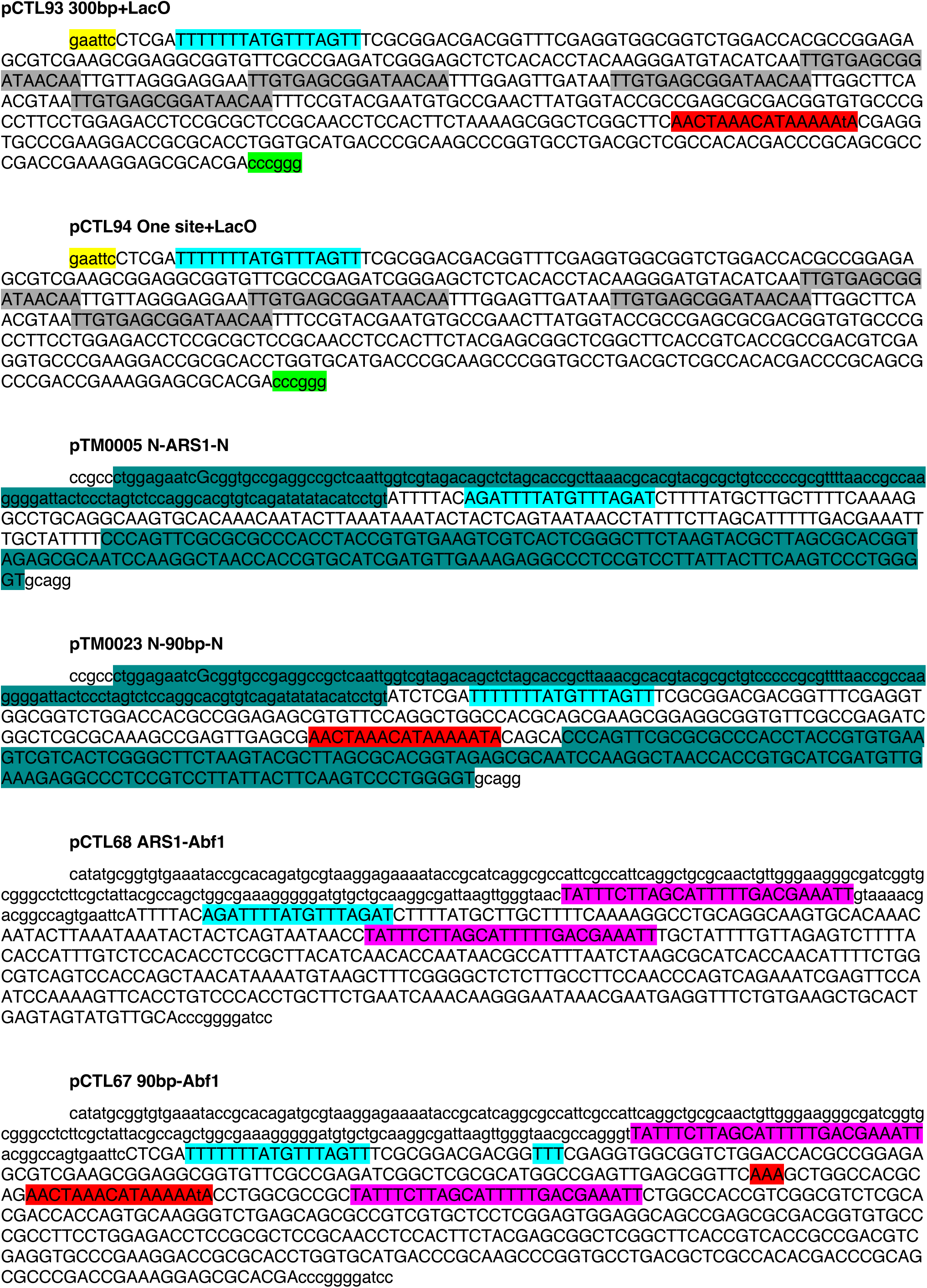
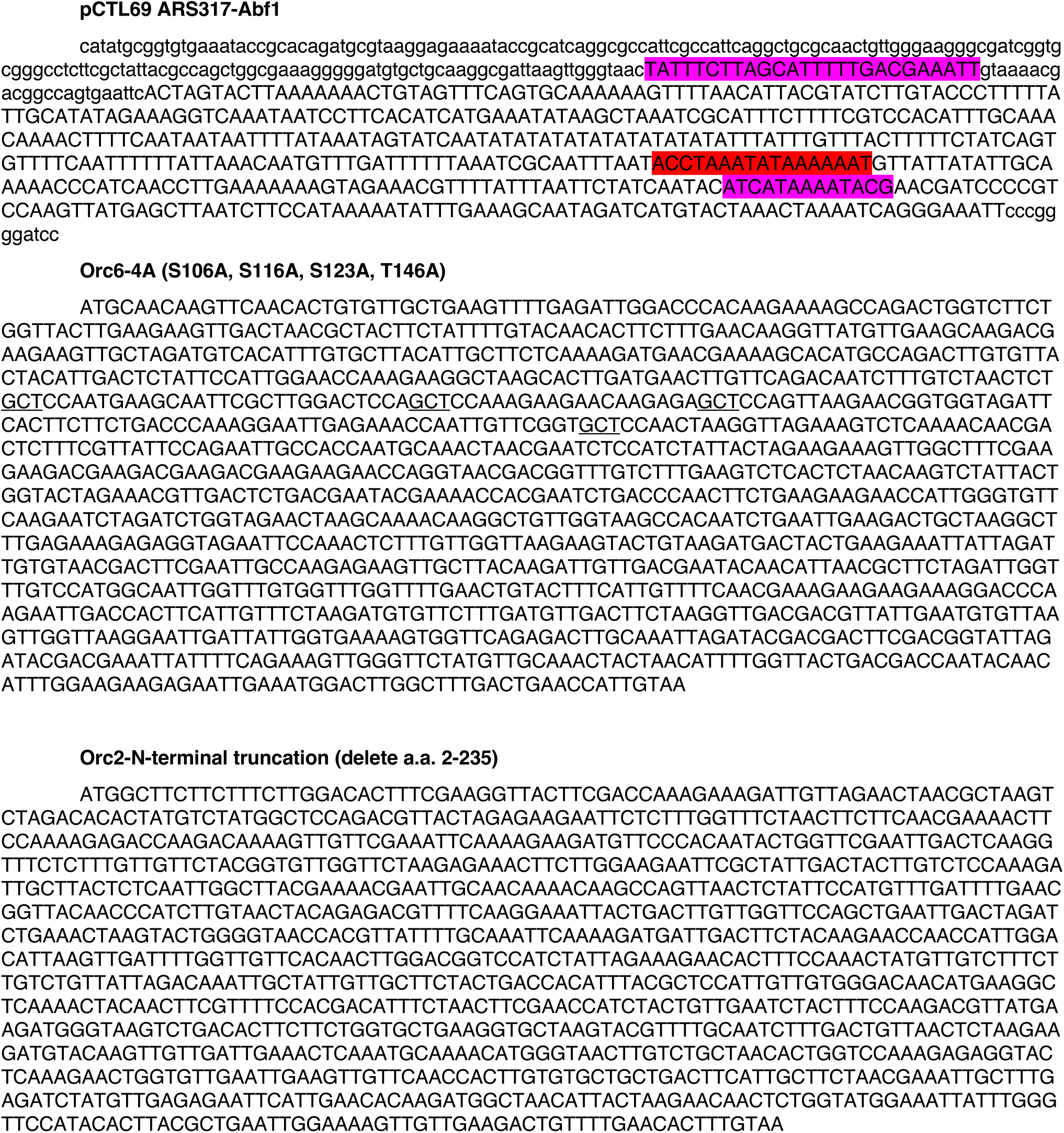

**Supplementary Table 1:**
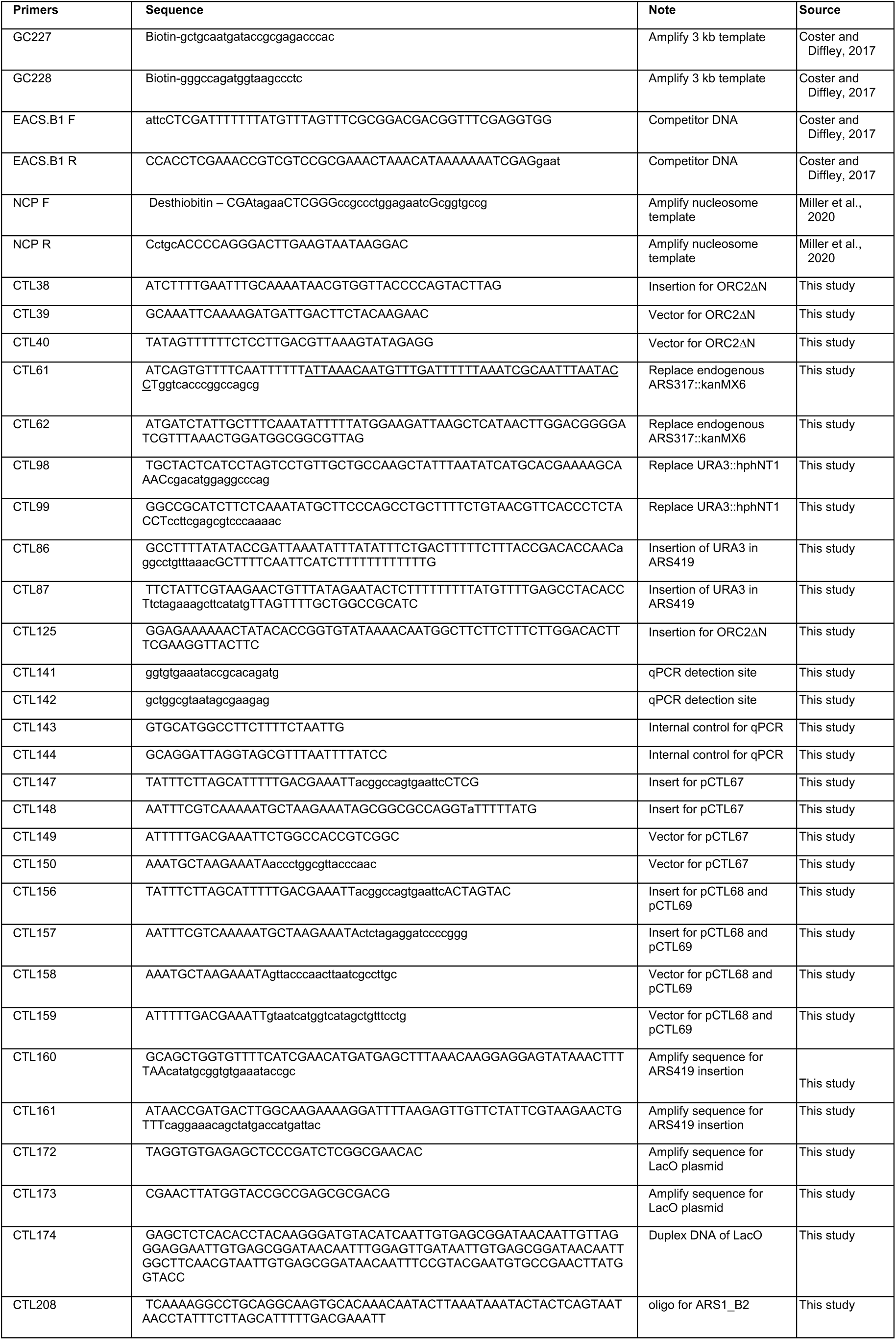
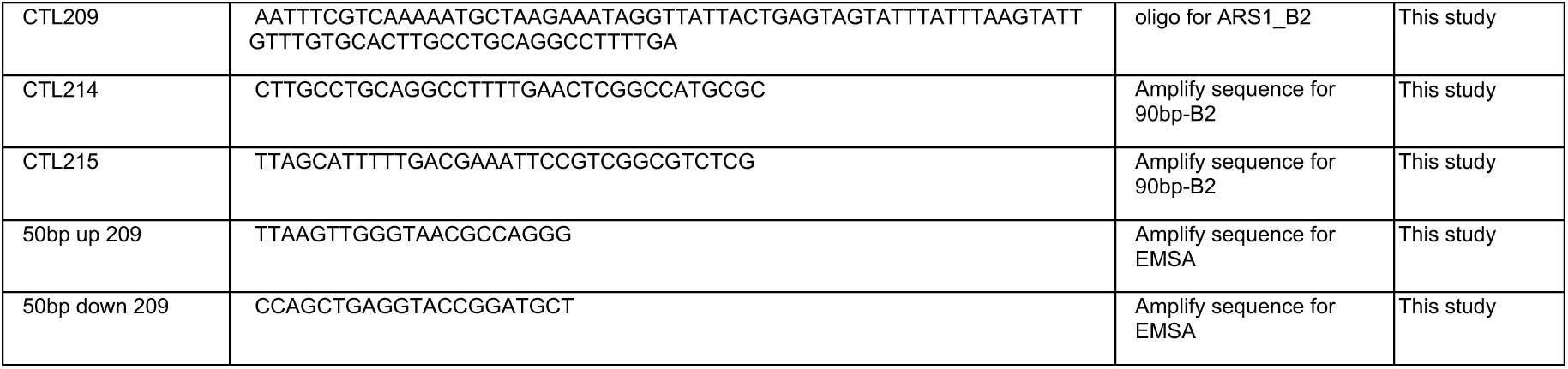
Primers.

**Supplementary Table 2:**
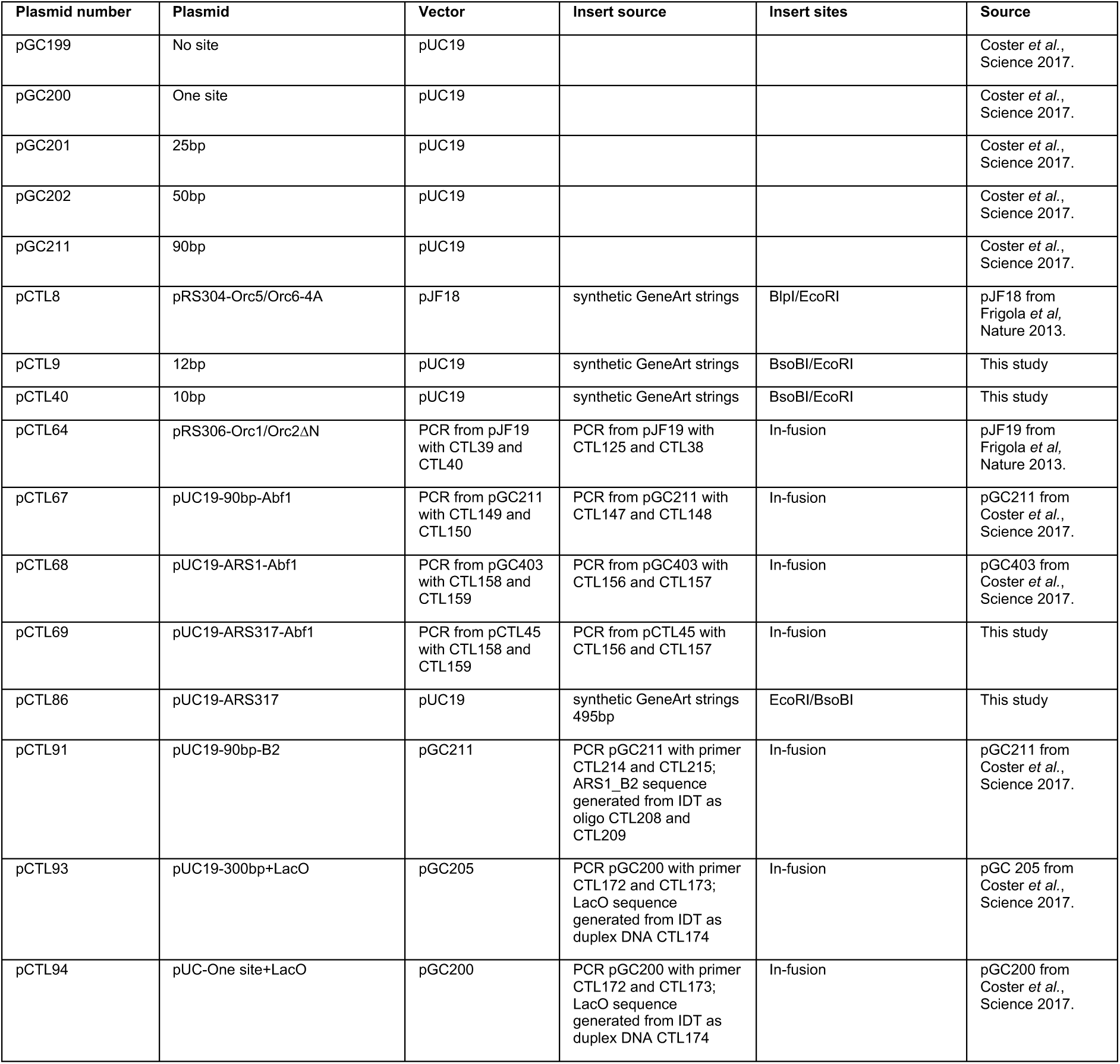
Plasmids.

**Supplementary Table 3:** Yeast strains.

| Yeast strain | Background | Genotype | Source |  |
| --- | --- | --- | --- | --- |
| ySDORC | W303-1a | *MATa ade2-1 ura3-1 his3-11,15 trp1-1 leu2-3,112 can1-100*<br>*bar1::Hyg*<br>*pep4::KanMx*<br>*his3::pRS303-Gal1,10-ORC3/4 (HIS3)*<br>*ura3::pRS306-Gal1,10-ORC1/2 CBP-Orc1 (URA3)*<br>*trp1::pRS304-Gal1,10-ORC5/6 (TRP1)*<br>*MATa ade2-1 ura3-1 his3-11,15 trp1-1 leu2-3,112 can1-100*<br>*bar1::Hyg*<br>*pep4::KanMx* | Frigola *et al.*, Nature 2013. |  |
| YGC229 | W303-1a | *his3::pRS303-Gal1,10-Cdt1/Gal4 (HIS3)*<br>*ura3::pRS306-Gal1,10-Mcm2/3 (URA3) CBP-TEV-NanoLuc-Mcm3*<br>*trp1::pRS304-Gal1,10-Mcm4/5 (TRP1)*<br>*leu2::pRS305-Gal1,10-Mcm6/7 (LEU2)*<br>*MATa ade2-1 ura3-1 his3-11,15 trp1-1 leu2-3,112 can1-100*<br>*bar1::Hyg*<br>*pep4::KanMx* | Coster *et al.*, Science 2017. |  |
| yCTL6<br>(Orc6-4A) | W303-1a | *MATa ade2-1 ura3-1 his3-11,15 trp1-1 leu2-3,112 can1-100*<br>*bar1::Hyg*<br>*pep4::KanMX*<br>Flag at WT ORC6 (Leu) c-terminal<br>*ura::URApRS306/SDORC1,2*<br>*his::HISpRS303/SDORC3,4*<br>*trp::TRPpRS304/SDORC5,ORC6-4S/T-A* | This study |  |
| yCTL17<br>(Orc2_deltaN) | W303-1a | *MATa ade2-1 ura3-1 his3-11,15 trp1-1 leu2-3,112 can1-100*<br>*bar1::Hyg*<br>*pep4::KanMX*<br>Flag at WT ORC2 (Nat) C-terminal<br>*ura::URApRS306/SDORC1,ORC2_del2-235*<br>*his::HISpRS303/SDORC3,4*<br>*trp::TRPpRS304/SDORC5,ORC6* | This study | RE digestion with NcoI on pCTL64 |
| yCTL28<br>(ARS1Δ) | A364a | *MATa ade2 ade3 ura3-52 trp1-289 leu2-3,112*<br>*bar1::LEU2*<br>*ORC2*<br>*ORC6*<br>*ura3-52::GAL1p-Δntcdc6, URA3 ΔURA3::hph*<br>*cdc20::MET3p-HA3-CDC20, TRP1*<br>*MCM7-2NLS*<br>*ARS317::KanMX6 ΔARS419::ARS1Δ* | This study | PCR on pGC404 with CTL160 and CTL161.<br>pGC404 from Coster *et al.*, Science 2017. |
| yCTL29<br>(ARS1) | A364a | *MATa ade2 ade3 ura3-52 trp1-289 leu2-3,112*<br>*bar1::LEU2*<br>*ORC2*<br>*ORC6*<br>*ura3-52::GAL1p-Δntcdc6, URA3 ΔURA3::hph*<br>*cdc20::MET3p-HA3-CDC20, TRP1*<br>*MCM7-2NLS*<br>*ARS317::KanMX6 ΔARS419::ARS1* | This study | PCR on pCTL68 with CTL160 and CTL161 |
| yCTL30<br>(ARS317) | A364a | *MATa ade2 ade3 ura3-52 trp1-289 leu2-3,112*<br>*bar1::LEU2*<br>*ORC2*<br>*ORC6*<br>*ura3-52::GAL1p-Δntcdc6, URA3 ΔURA3::hph*<br>*cdc20::MET3p-HA3-CDC20, TRP1*<br>*MCM7-2NLS*<br>*ARS317::KanMX6 ΔARS419::ARS317* | This study | PCR on pCTL69 with CTL160 and CTL161 |
| yCTL33<br>(90bp) | A364a | *MATa ade2 ade3 ura3-52 trp1-289 leu2-3,112*<br>*bar1::LEU2*<br>*ORC2*<br>*ORC6*<br>*ura3-52::GAL1p-Δntcdc6, URA3 ΔURA3::hph*<br>*cdc20::MET3p-HA3-CDC20, TRP1*<br>*MCM7-2NLS*<br>*ARS317::KanMX6 ΔARS419::90bp* | This study | PCR on pCTL67 with CTL160 and CTL161 |

## Notes

### Competing Interest Statement

The authors have declared no competing interest.

